# Transcriptional overlap links DNA hypomethylation with DNA hypermethylation at adjacent promoters in cancer

**DOI:** 10.1101/2021.05.21.445142

**Authors:** Jean Fain, Axelle Loriot, Anna Diacofotaki, Aurélie Van Tongelen, Charles De Smet

## Abstract

DNA methylation is an epigenetic mark associated with gene repression. It is now well established that tumor development involves alterations in DNA methylation patterns, which include both gains (hypermethylation) and losses (hypomethylation) of methylation marks in different genomic regions. The mechanisms underlying these two opposite, yet co-existing, alterations in tumors remain unclear. While studying the human *MAGEA6*/*GABRA3* gene locus, we observed that DNA hypomethylation in tumor cells can lead to the activation of a long transcript (*CT-GABRA3*) that overlaps downstream promoters (*GABRQ* and *GABRA3*) and triggers their hypermethylation. Overlapped promoters displayed increases in H3K36me3, a histone mark known to be deposited during progression of the transcription machinery and to stimulate de novo DNA methylation. Consistent with such a processive mechanism, increases in H3K36me3 and DNA methylation were observed over the entire region covered by the *CT-GABRA3* overlapping transcript. Importantly, experimental induction of *CT-GABRA3* by depletion of DNMT1 DNA methyltransferase, resulted in a similar pattern of increased DNA methylation in the *MAGEA6*/*GABRA3* locus. Bioinformatics analyses in lung cancer datasets identified other genomic loci displaying this process of coupled DNA hypo- and hypermethylation. In several of these loci, DNA hypermethylation affected tumor suppressor genes, e.g. *RERG* and *PTPRO*. Together, our work reveals that focal DNA hypomethylation in tumors can indirectly contribute to hypermethylation of nearby promoters through activation of overlapping transcription, and establishes therefore an unsuspected connection between these two opposite epigenetic alterations.

## INTRODUCTION

Cancer development is driven in part by the accumulation of epigenetic alterations, which render chromatin permissive to changes in gene expression patterns. As a result, tumor cells acquire increased plasticity, thereby facilitating their evolution towards full malignancy [1]. Epigenetic alterations concern in particular DNA methylation, a chemical modification of cytosines in CpG sequences that is associated with long-term transcriptional repression [2]. DNA methylation changes in tumors include gains (hypermethylation) within normally unmethylated gene promoters, and at the same time losses (hypomethylation) in other genomic sequences [3]. The mechanisms underlying these contrasting changes in DNA methylation patterns in tumors are only partially elucidated, and evidence so far suggest that DNA hypermethylation and hypomethylation result from two independent processes [4, 5].

DNA hypermethylation has a well-established role in tumor progression, as it can lead to irreversible silencing of genes with tumor suppressor functions [6]. DNA hypomethylation on the other hand, appears to promote tumor development by increasing genomic instability, and by inducing ectopic activation of genes with oncogenic functions [7]. Many of these latter genes belong to the class of so-called “cancer-germline” (CG) genes, as their expression in healthy adults is normally restricted to testicular germ cells [8]. It has indeed been demonstrated that CG genes rely primarily on DNA methylation for repression in non-expressing cells, and that DNA demethylation is a sufficient trigger for their activation in a variety of tumors [9-11]. Evidence has accumulated indicating that some CG genes contribute to tumor progression, notably by encoding proteins that regulate processes of cell proliferation, death resistance, metabolic adaptation, and DNA repair [12, 13].

Recently, we discovered a novel CG transcript (*CT-GABRA3*) showing DNA hypomethylation-dependent activation in a variety of tumors, including melanoma and lung cancer [14, 15]. The *CT-GABRA3* transcript is non-coding, extends over a large distance (530 kb), and overlaps the Gamma-Aminobutyric Acid Type A Receptor Subunit Alpha3 (*GABRA3*) gene, starting ∼250 kb downstream. Of note, *CT-GABRA3* shares a bidirectional promoter with the well described *MAGEA6* CG gene, and both transcripts are most often co-expressed. An intriguing observation was that in the melanoma cell lines where we detected hypomethylation and activation of *MAGEA6*/*CT-GABRA3*, the promoter of *GABRA3* exhibited marked hypermethylation. This suggested that promoter hypomethylation and subsequent transcriptional activation of *CT-GABRA3* might trigger DNA hypermethylation of the downstream *GABRA3* overlapped promoter. Epigenetic processes involving overlapping transcription have indeed been implicated in the establishment of DNA methylation marks at parentally imprinted sites, and in intragenic promoters during development [16, 17]. The underlying mechanism involves deposition of H3K36me3 modification along with the transcriptional machinery, and consequent recruitment of *de novo* DNA methyltransferases. This mechanism also explains the fact that actively transcribed genes usually show higher CpG methylation within their body, probably as a way to prevent spurious transcription initiation downstream of the transcription start site [18-20].

In the present study, we set out to validate the relationship between the methylation status of *CT-GABRA3* and *GABRA3* promoters in tumors. The involvement of transcriptional overlap in this epigenetic coupling was investigated. We then used a bioinformatics approach to explore the possibility that this process of coupled hypo/hypermethylation of CpGs occurs in other parts of the genome of cancer cells, and contributes to DNA hypermethylation of tumor suppressor genes.

## MATERIAL AND METHODS

### Cell culture

All human melanoma (MZ2-MEL, Mi13443, LB39-MEL, LB2667-MEL, Mi1811) and lung cancer (SKMES-1, GLCP1, LB37, NCI-H661) cell lines were obtained from the Brussels branch of the Ludwig Institute for Cancer Research. Melanoma cell lines were cultured as previously described [21]. SKMES1, GLCP1 and LB37 cell lines were cultured in IMDM (Life Technologies) and NCI-H661 was cultured in RPMI (Life Technologies) medium, supplemented with 10% of fetal bovine serum (FBS, Sigma), 1x of non-essential amino acids (Life Technologies) and 1x penicillin/streptomycin (Life Technologies). Early passage human normal epidermal melanocytes were received from E. De Plaen (Ludwig Institute for Cancer Research, Belgium) and were cultured in Ham’s F10 medium (Life Sciences) supplemented with 6 mM Hepes, 1 x MelanoMax supplement (Gentaur), and 10% FBS. For 5-azadC induction experiments, cells grown to 60-70% confluency were exposed to a single dose of 2μM 5-aza-2ʹ -deoxycytidine (Sigma-Aldrich) diluted in 1:1 acetic acid/water. Treated cells were maintained in culture during 6 days before RNA extraction.

### RT-PCR and qPCR analyses

RNA of tissue samples was purchased from Ambion (Life Technologies). RNA of cell lines was extracted using TriPure Isolation Reagent (Roche Diagnostics GmbH). Reverse transcription was performed on 2 μg of total RNA using M-MLV Reverse transcriptase and random hexamers (Invitrogen). For PCR reactions, we used the DreamTaq Kit (Thermo Fisher Scientific), incorporating 1/40 of the reverse transcription mixture in a final reaction volume of 20 μl. PCR reactions were visualized after electrophoresis in an ethidium bromide-stained agarose gel. For qPCR reactions, we used KAPA SYBR FAST (Sigma-Aldrich), incorporating 1/40 of the reverse transcription mixture in a final reaction volume of 10μl. All reactions were carried out according to the manufacturer’s instructions. All primers are listed in the supplementary table S2.

### Sodium bisulfite sequencing

Sodium bisulfite genomic sequencing of *CT-GABRA3* and *GABRA3* promoter regions was performed as previously described (48). Primer used for nested PCR amplification of bisulfite treated DNA are listed in the supplementary table S2.

### Processing of public RNA-seq raw data

Fastq files of the 26 LUAD cell lines were downloaded from the DNA Data Bank of Japan (PRJDB2256). Fastq files of normal lung and testis tissues were downloaded from Sequence Read Archive of NCBI (PRJNA34535 & PRJEB4337). All accession numbers are listed in supplementary table S3. Read alignment, *de novo* transcriptome assembly, and quantification of full-length referenced and unreferenced transcripts, were performed as described in the supplementary methods. For calculation of *CT-GABRA3* and *CT-RERG* expression levels in LUAD cell lines and normal tissues, transcripts originating from the same TSS were summed. LUAD cell lines were considered positive for *CT-GABRA3* or *CT-RERG* expression when the corresponding transcript showed a TPM ≥ 1.

### DNA methylation analyses in sperm, lung and LUAD cell lines

#### 1) Data collection

Whole genome bisulfite sequencing (WGBS) data for sperm and lung are provided by the NIH Roadmap epigenomics [22]. Normalized hg19 WGBS-seq data for sperm and lung were downloaded through the NCBI Gene Expression Omnibus, and were converted to hg38 using liftOver v1.10.0 R package. As corresponding data processing workflow does not allow multi-mapping of reads, methylation data for duplicated genomic regions, such as that containing the *CT-GABRA3*/*MAGEA6* promoter [14], were not available for sperm and lung. For LUAD cell lines, normalized hg38 target-captured bisulfite sequencing (Methyl-seq) data were downloaded from DBTSS v9 [23]. Only a fraction of genomic CpGs (∼12%) are covered by the Methyl-seq method. Of note, three LUAD cell lines displaying ambiguous expression and DNA methylation results for the highly similar *MAGEA6* and *MAGEA3* genes [14] were ignored for the analysis of the *CT-GABRA3/MAGEA6* promoter methylation status. All accession numbers are listed in supplementary table S3. *2) Data analyses:* For regional DNA hypermethylation analysis, we studied the methylation status of all CpGs located between +1 kb of the TSS and up to the 5’ end of the OTr (= region B: 530 kb for *CT-GABRA3* and 240 kb for *CT-RERG*). Genomic segments of the same size were used to investigate CpG methylation levels in neighboring regions (regions A and C). For the upstream region A, CpGs located within 1kb upstream from the TSS of the OTr were excluded from the analysis. For analyses in LUAD cell lines, we only retained CpGs for which the methylation status could be determined in >70% of the cell lines.

### ChIP-seq data collection and analysis

Hg38 ChIP-seq data for H3ac, H3K4me3, H3K9me3, H3K27me3, H3K36me3 histone marks (and input) of the 26 LUAD cell lines were downloaded from DBTSS v9. To quantify histone modifications within promoter (TSS +/-1kb) or genomic regions of interest in LUAD cell lines, we computed the total number of reads mapped to the corresponding genomic segment, divided by the sum of all reads generated in the same experiment, and multiplied by 10^6^ to obtain Reads Per Million (RPM) values.

### TCGA consortium datasets

#### 1) Data collection

Normalized hg19 RNA-seq data with exon-level quantification and Infinium Human Methylation 450K datasets for skin cutaneous melanoma (SKCM), lung adenocarcinoma (LUAD), liver hepatocellular carcinoma (LIHC), and kidney papillary cell carcinoma (KIRP) were downloaded from The Cancer Genome Atlas (TCGA) consortium [24], using TCGAbiolinks v2.14.1 R-package [25]. Hg19 coordinates were converted to hg38 using liftOver v1.10.0 R package. Only unique primary (−01A) and metastatic (−06A) tumor samples, as well as unique normal adjacent tissues (−11A), for which both RNA-seq and Infinium methylation data were available, were analyzed. RNA-seq exon expression levels are expressed as Reads Per Kilobase per Million (RPKM). *2) Expression analyses:* Since *CT-GABRA3* and *CT-RERG* transcripts variants are not annotated in TCGA-derived datasets, we resorted to exon quantification to determine their expression status. Presence or absence of the canonical exon 1 allowed to distinguish *CT-GABRA3* or *CT-RERG* transcript variants versus *GABRA3* or *RERG* referenced transcripts, respectively. Thresholds were determined as follows: samples were considered positive for *CT-GABRA3* expression if exon 1 displayed ≤ 0.1 RPKM and exon 2 ≥ 1 RPKM; *CT-RERG* expression was positive when exon 1 displayed ≤ 0.4 RPKM and exon 5 ≥ 1 RPKM. *3) DNA methylation analyses:* For regional DNA hypermethylation analyses, Infinium methylation levels (beta values) were examined for all CpG probes located in regions A, B and C regions, demarcated as described here above.

### DNMT1 depletion experiments of O’Neill’s study

Illumina HumanHT-12 V4.0 expression data (GSE90012) and Infinium Human Methylation 450K data (GSE90011) were downloaded from NCBI Gene Expression Omnibus database. The following probes were used for expression analysis of the genes of interest: *DNMT1* (ILMN_1715551), *MAGEA6* (ILMN_2372681), and (*CT-*)*GABRA3* (ILMN_1715551). For regional hypermethylation analyses, Infinium methylation levels (beta values) were examined for all CpG probes located in regions A, B and C regions, demarcated as described here above.

### Bioinformatics workflow for the identification of overlapped promoter hypermethylation

Identification of genomic loci that harbor an activated transcript leading to hypermethylation of a downstream promoter in LUAD cell lines, was performed by using a pipeline conducted in Perl programming language (scripts are available upon request), and applied to RNA-seq and Methyl-seq data of LUAD cell lines (DNA Data Bank of Japan, DBTSS), as well as RNA-seq and WGBS data of normal lung (Sequence Read Archive of NCBI, ENCODE). Initial processing of these RNA-seq datasets was described above. Details on the procedures to select transcripts activated in LUAD cell lines and potential overlapped promoters, and to establish correlations between overlapping transcript expression and overlapped promoter methylation, are described in the supplementary methods.

### Statistical analysis and graphical representations

Statistical analysis was computed in R v3.6.1 (http://www.R-project.org). Graphs and heatmaps were generated using R packages ggplot2 (v3.3.2) and ComplexHeatmap (v2.2.0). Benjamini-Hochberg correction was used for adjustment of *p*-values.

## RESULTS

### *MAGEA6/CT-GABRA3* promoter hypomethylation correlates with *GABRA3* promoter hypermethylation in melanoma

Recently, our studies focused on the human *MAGEA6*/*GABRA3* gene locus on chromosome X [14, 15]. We demonstrated the existence of a bidirectional promoter (*MAGEA6*/*CT-GABRA3*) driving expression of both *MAGEA6* and *CT-GABRA3* transcripts in testis. *CT-GABRA3* starts ∼250-kb upstream of another gene (*GABRA3*), comprises several specific exons in its 5’ part, and then overlaps *GABRA3*, of which it comprises all but exon 1 (Fig. 1A). *MAGEA6* and *CT-GABRA3* become aberrantly co-activated in a significant proportion of tumors, including melanoma, and our previous studies indicated that this was caused by DNA demethylation of the *MAGEA6*/*CT-GABRA3* promoter [14, 15]. Evidence for this was in part provided by bisulfite sequencing results, as shown in figure 1B, showing that the *MAGEA6*/*CT-GABRA3* promoter is initially methylated in normal melanocytes, the cell of origin of melanoma. Contrastingly, in melanoma cells where *MAGEA6* and *CT-GABRA3* are activated, the promoter is completely demethylated, including at critical CpG sites located near the transcription start sites (Fig. 1B). We showed previously that methylated CpGs located near the TSS are most critical for transcriptional repression of CG genes [9]. In the present study, we analyzed the methylation status of the *GABRA3* promoter, which normally remains poorly methylated in all tissues (supplementary Fig.S1). Intriguingly, DNA hypermethylation of the *GABRA3* promoter was observed specifically in the melanoma cells where the overlapping *CT-GABRA3* transcript was produced (Fig. 1B).

**Figure 1.**
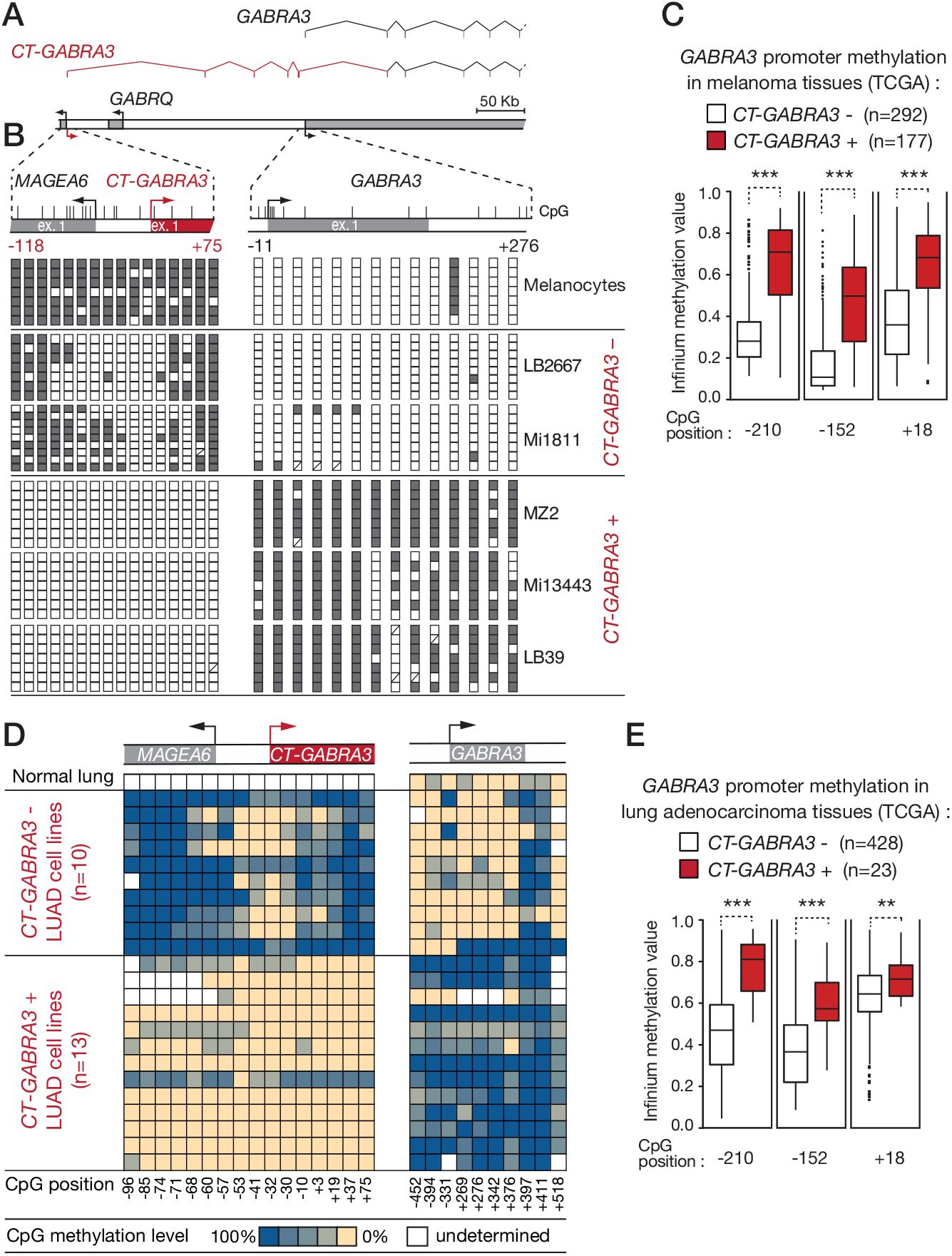
DNA hypomethylation of the *MAGEA6*/*CT-GABRA3* promoter correlates with DNA hypermethylation of the *GABRA3* promoter in melanoma and lung adenocarcinoma. (*A*) Schematic representation of the *GABRA3* locus, with broken arrows indicating transcription start sites. The exon/intron structure of *CT-GABRA3* and *GABRA3* transcript variants is shown above. (*B*) Clonal bisulfite sequencing of *MAGEA6*/*CT-GABRA3* and *GABRA3* 5’-regions. Vertical bars indicate location of CpG sites with positions relative to the transcription start site. Open and filled squares represent unmethylated and methylated CpG sites, respectively, and each row represents a single clone. *CT-GABRA3* expression status (+) or (−) in melanoma cell lines is indicated. (*C*) Melanoma tissue samples from the TCGA were grouped according to *CT-GABRA3* expression status (inferred from RNA-seq data), and the methylation level of three CpG sites embedded within the *GABRA3* 5’-region (position relative to TSS) were determined through the analysis of Infinium methylation data (probe intensity ratio). *** Welch’s t-test, adjusted *p*-value <0.001 (*D*) Methylation level of CpG sites within the *MAGEA6*/*CT-GABRA3* and *GABRA3* 5’-regions in lung adenocarcinoma (LUAD) cell lines (CpG positions are expressed relative to the TSS). Methylation levels were calculated on the basis of Methyl-seq data from the DBTSS database. *CT-GABRA3* expression status in LUAD cell lines was inferred from RNA-seq data. (*E*) The same analysis as described in *D* was applied to lung adenocarcinoma samples from the TCGA. ** and *** Mann-Whitney test, adjusted *p*-value <0.01 and <0.001, respectively.

To further extend this observation, we searched to establish a similar relationship in a large number of melanoma tissue samples. Interestingly, examination of transcriptomic (RNA-seq) and methylomic (Infinium methylation assay) data from the Cancer Genome Atlas (TCGA), revealed a significant association between expression of *CT-GABRA3* and hypermethylation of the downstream *GABRA3* promoter in melanoma (Fig. 1C). Of note, this association was not affected by gender, thereby excluding a process related to X chromosome inactivation (supplementary Fig.S2).

### Concurrent *CT-GABRA3* activation and *GABRA3* hypermethylation in lung adenocarcinoma

*CT-GABRA3* is expressed not only in melanoma, but also in lung cancer. We decided to verify the association between hypomethylation/activation of *CT-GABRA3* and hypermethylation of *GABRA3* in this tumor type. To this end, we first exploited the Database of Transcription Start Sites (DBTSS), which contains various multi-omics data for a set of human lung adenocarcinoma (LUAD) cell lines [23]. Twenty three cell lines were grouped according to the status of expression of *CT-GABRA3*, which was defined on the basis of RNA-seq data [14]. Among the 23 LUAD cell lines, 13 (57%) scored positive for *CT-GABRA3* (as well as *MAGEA6*) activation (Fig. 1D). Using Methyl-seq datasets, we then evaluated the DNA methylation levels of *MAGEA6*/*CT-GABRA3* and *GABRA3* promoters in the different LUAD cell lines. The results revealed that *CT-GABRA3* activation in these cell lines is associated with hypomethylation of its promoter, as expected, but also with hypermethylation of the promoter region of *GABRA3* (Fig. 1D). To verify if this association also pertains *in vivo* in lung adenocarcinoma tissue samples, we resorted to the analysis of TCGA datasets (Fig. 1E). This confirmed significant association between *CT-GABRA3* activation and *GABRA3* promoter hypermethylation in lung adenocarcinoma.

### DNA hypermethylation extends all over the *CT-GABRA3* transcription unit

The 530 kb genomic segment covered by the *CT-GABRA3* transcript variant comprises another gene with brain-specific expression, *GABRQ* (Fig 2A). Unlike *GABRA3, GABRQ* is normally transcribed in the opposite direction to *CT-GABRA3*. Examination of Methyl-seq data from the DBTSS database revealed that CpGs within the *GABRQ* promoter also displayed increased methylation in LUAD cell lines that express *CT-GABRA3* (Fig. 2A). A similar observation was made for most other CpGs assessed within the 530 kb locus, thereby suggesting that *CT-GABRA3* transcription exerts a regional effect of DNA hypermethylation (Fig. 2A). To evaluate the extent of this effect, changes of CpG methylation levels were analyzed in regions located immediately upstream (region A) and downstream (region C) of the locus defined by the *CT-GABRA3* transcription unit (region B; Fig. 2B). Methyl-seq data in LUAD cell lines indicated that whereas *CT-GABRA3* activation was associated with DNA hypermethylation within its transcription unit (region B), it was instead associated with reduced DNA methylation levels in the neighboring regions A and C. This likely reflects the fact that activation of *CT-GABRA3* occurs preferentially in tumor cells with global genome hypomethylation (Fig. 2C). Analysis of TCGA methylomic datasets confirmed the existence of a similar profile of regional DNA hypermethylation, limited to the *CT-GABRA3* transcription unit, in melanoma and lung adenocarcinoma tissue samples (Fig. 2D).

**Figure 2.**
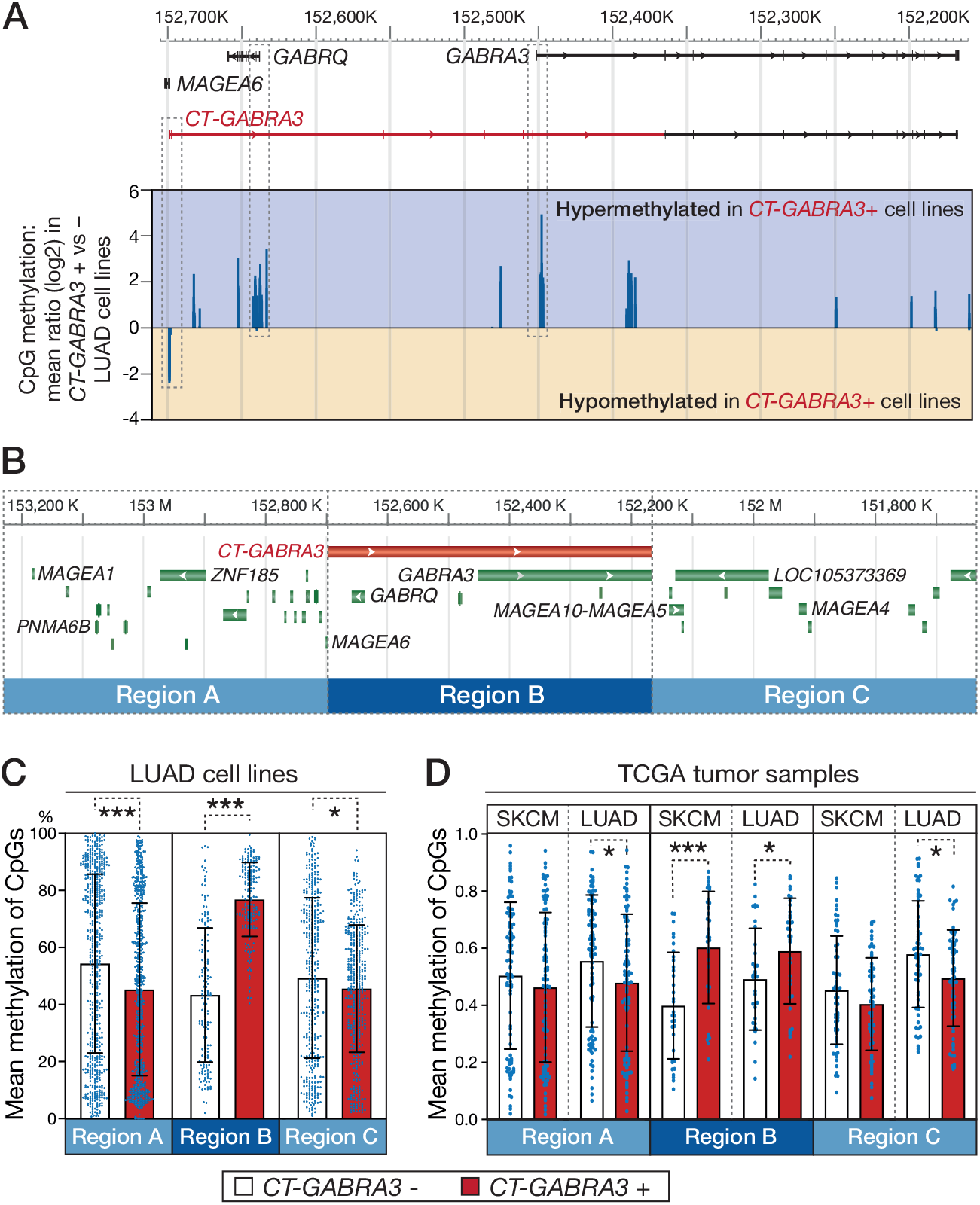
*CT-GABRA3* expression correlates with hypermethylation of CpGs embedded within its entire transcription unit. (*A*) Using Methyl-seq data from DBTSS, CpG methylation ratios between *CT-GABRA3*-positive and -negative LUAD cell lines were calculated. Values for all available CpGs within the *GABRA3* locus are represented (log2). Positive or negative values indicate hypermethylation or hypomethylation, respectively, of the CpG in *CT-GABRA3*-positive versus -negative LUAD cell lines. Hypermethylation includes CpGs within the promoter of *GABRQ*. (*B*) Schematic representation of the three defined 530 kb genomic segments, corresponding to the *CT-GABRA3* transcription unit (Region B), and the two neighboring segments (Regions A and C). (*C*) LUAD cell lines were divided in two groups, according to *CT-GABRA3* expression status, and Methyl-seq datasets from DBTSS were used to determine mean methylation levels (% methylation) of all CpGs contained in each of the three genomic regions defined in *B*. (*D*) A similar analysis, based on Infinium methylation data, was performed in melanoma (SKCM) and lung adenocarcinoma (LUAD) tissue samples from TCGA. * and *** Welch’s t-test, *p*-value <0.05 and <0.001, respectively.

### Experimental evidence demonstrating that DNA hypomethylation/activation of *CT-GABRA3* induces *de novo* methylation of overlapped CpGs

So far, our observations linking hypomethylation/activation of *CT-GABRA3* and hypermethylation of CpG sites located within its 530 kb transcription unit were only correlative. To establish a direct link between these two events, we explored experimental results obtained by O’Neill and colleagues [26], who generated three immortalized human fibroblast clones (of male origin) in which DNMT1 DNA methyltransferase was depleted following stable transfection of a specific shRNA vector (Fig. 3A). Interestingly, the authors reported that experimental depletion of DNMT1 resulted not only in losses, but also in gains of DNA methylation [26]. We decided to explore O’Neill’s datasets to find out the changes that occurred within the *MAGEA6/GABRA3* locus. Previous reports demonstrated that DNMT1 plays a key role in maintaining silencing of CG genes [21, 27, 28]. Consistently, cDNA microarray data revealed concurrent up-regulation of *MAGEA6* and *(CT-)GABRA3* (microarray probes do not distinguish *CT-GABRA3* and *GABRA3* variants) in DNMT1-depleted cell clones (Fig. 3B). We then analyzed Infinium methylation assay datasets generated for the different groups of cells to evaluate methylation levels of CpGs located in either the *CT-GABRA3* transcription unit (region B) or in the neighboring regions (region A and C, see Fig. 2B). The results revealed that, compared with the control cell line, all three DNMT1-depleted cell clones displayed significant increases of CpG methylation within the *CT-GABRA3* transcription unit (region B, Fig. 3C). Methylation levels of CpGs located in adjacent regions A and C, remained instead unchanged. Together these results demonstrate that hypomethylation/activation of *CT-GABRA3* is linked with a process of *de novo* methylation of CpGs located within its transcription unit.

**Figure 3.**
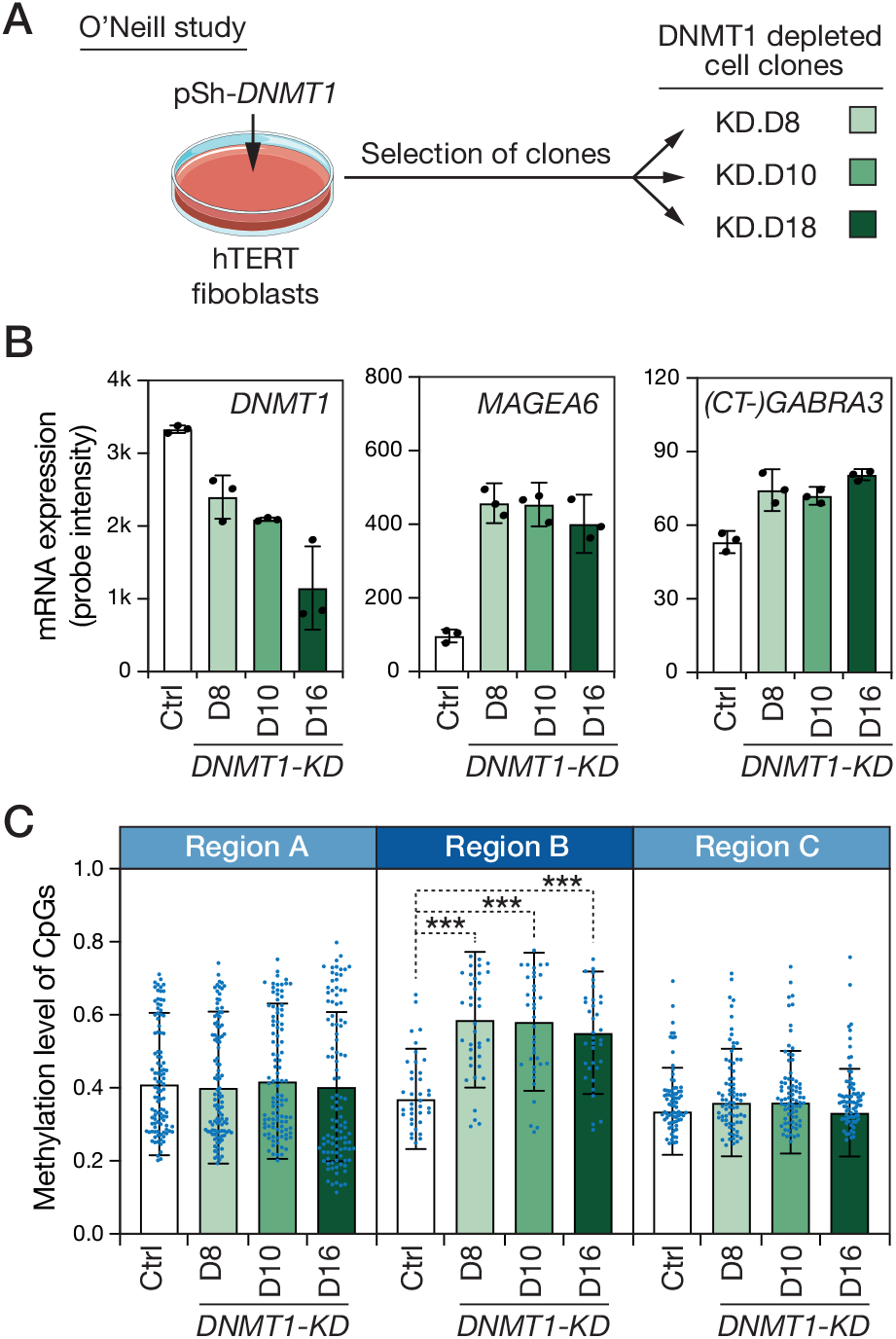
Experimental hypomethylation/activation of *CT-GABRA3* induces *de novo* methylation of CpGs within its transcription unit. (*A*) Diagram depicting O’Neill’s experiment. Clones were derived from hTERT-immortalized human male fibroblasts transfected with an anti-DNMT1 shRNA producing vector (pSh-*DNMT1*). (*B*) cDNA microarray data were analyzed to determine relative mRNA levels of *DMNT1, MAGEA6* and (*CT-*)*GABRA3* (the two transcript variants cannot be distinguished by microarray), in the control hTERT fibroblast cell line (Ctrl) and in the three DNMT1-depleted clones (D8, D10 and D18). (*C*) Infinium methylation data were used to determine mean methylation levels of all CpGs (mean probe intensity ratios) in the control cell line and the DNMT1-depleted clones. Analyses are provided for the *CT-GABRA3* transcription unit (Region B), and the two neighboring segments (Regions A and C, see Fig.2B). *** One-way paired ANOVA test with Dunnett’s correction, *p*-value <0.001.

### *CT-GABRA3* transcription in LUAD cells correlates with regional increases in H3K36me3

We next searched to determine if *CT-GABRA3* transcription also modifies histone marks within the overlapped genomic region. To this end, we analyzed ChIP-seq datasets of LUAD cell lines via the DBTSS platform, in order to evaluate the level of various histone marks around the transcription start site of *GABRA3*. These analyses revealed that *CT-GABRA3* transcription in LUAD cells correlates with a decrease in H3K27me3, which is consistent with initial presence of this mark, and its loss upon DNA hypermethylation through the previously described process of epigenetic switch (supplementary Fig. S3, and [29]). Strikingly, an enrichment in H3K36me3 within the 5’-region of *GABRA3* was also observed (Fig. 4A). Other histone marks, including H3K4me3, H3K9me3, and H3ac remained unchanged. Similar observations were made for the *GABRQ* promoter (supplementary Fig. S4).

**Figure 4.**
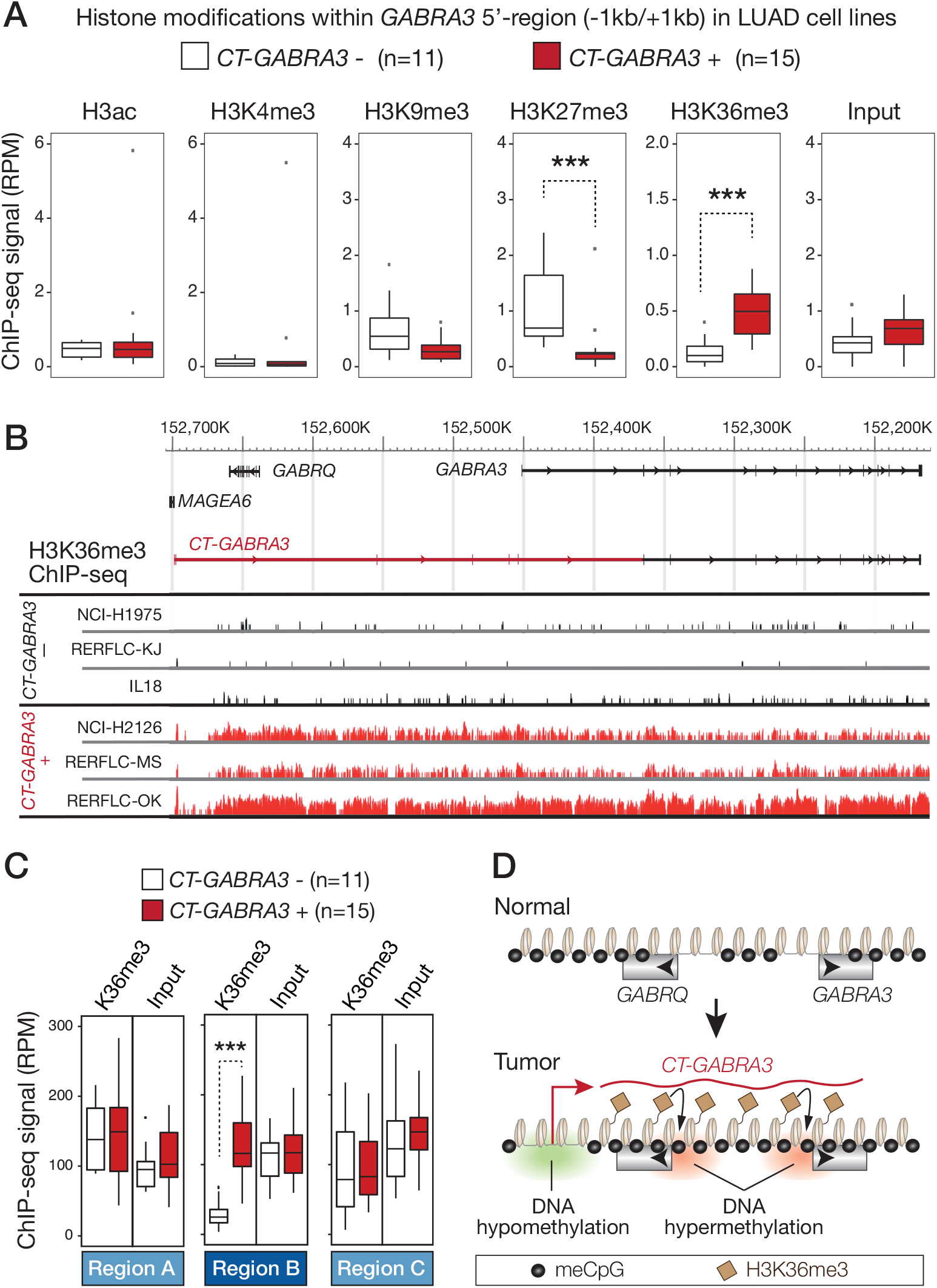
*CT-GABRA3* transcription correlates with regional increases in H3K36 trimethylation. (*A*) ChIP-seq results for the indicated histone modifications were extracted from the DBTSS platform, and mean amounts of reads corresponding to the *GABRA3* 5’ region (−1b/+1kb) were compared in LUAD cell lines that do or do not express *CT-GABRA3*. *** Mann-Whitney test, adjusted *p*-value <0.001. (*B*) ChIP-seq profile for H3K36me3 (DBTSS browser) within the entire *CT-GABRA3* transcription unit, in 3 LUAD cell lines that do not express *CT-GABRA3* (−) and 3 that do express it (+). (*C*) Mean amounts of ChIP-seq reads for H3K36me3 or control input in the three defined 530 kb genomic segments (Region B: *CT-GABRA3* transcription unit; Regions A, C: neighboring segments), were compared in LUAD cell lines that do or do not express *CT-GABRA3*. *** Mann-Whitney test, *p*-value <0.001. (D) Model establishing the link between DNA hypomethylation/activation of *CT-GABRA3* and DNA hypermethylation of *GABRQ* and *GABRA3* promoters in tumor development.

H3K36me3 is classically enriched over the body of actively transcribed genes, as it is deposited along with the transcription machinery. It has been shown that H3K36me3 favors local DNA methylation by attracting DNMT3 methyltransferases [16, 18, 20]. Inspection of the distribution of H3K36me3 within the entire 530 kb genomic region covered by *CT-GABRA3* transcription, revealed that enrichment of this histone mark in *CT-GABRA3*-positive LUAD cell lines already becomes apparent 15 to 20-kb downstream of the transcription start site, and extends up into the *GABRA3* promoter and beyond (Fig. 4B). Examination of ChIP-seq signals in neighboring segments (regions A and C, see Fig. 2B), indicated that increases in H3K36me3 were limited to the region overlapped by *CT-GABRA3* transcription (region B, Fig. 4C). Together, these observations suggest that *CT-GABRA3* transcription is accompanied by deposition of the repressive H3K36me3 histone mark, and leads thereby to increased susceptibility of the entire transcription unit to DNA hypermethylation. This model explains how DNA hypomethylation, and concurrent transcriptional activation, can be connected with hypermethylation of adjacent promoters (Fig. 4D).

### Other gene promoters displaying DNA hypermethylation in association with overlapping transcription in lung adenocarcinoma cells

An important issue was to determine if genes other than *GABRA3* and *GABRQ*, and in particular tumor suppressor genes, rely on a similar process of DNA hypomethylation-induced overlapping transcription to become hypermethylated in tumors. To this end, we examined the RNA-seq and Methyl-seq data obtained from LUAD cell lines (DBTSS) by applying a computational selection procedure to identify genomic loci that displayed the following features: i) ectopic activation in at least one LUAD cell line of a transcript that is not expressed in normal lung, ii) the ectopic transcript overlaps one or several downstream promoter(s) in either sense or anti-sense orientation, iii) the downstream overlapped promoter(s) (OPr) is(are) unmethylated in normal lung, and its hypermethylation is correlated with activation of the overlapping transcript (OTr) (Fig. 5A). This led to a list of 35 genomic loci, besides that containing *GABRA3* and *GABRQ*. In three of these loci, activation of the overlapping transcript was correlated with DNA hypermethylation in not only one but two overlapped genes. Hence, our search identified 38 genes showing promoter hypermethylation in association with activation of an overlapping transcript in LUAD cell lines (Fig. 5B, supplementary Table S1). Overlapped promoters were located 2 kb to 128 kb downstream of the OTr transcription start site, in either sense or antisense orientation, and generally contained a high density of CpGs (Fig. 5B, C). Moreover, examination of ChIP-seq data revealed that 87% of the overlapped promoters displayed significant enrichment of H3K36me3 in the LUAD cell lines that express the corresponding overlapping transcript (Pearson correlation coefficient >0.5, adjusted *p-*value <0.05; Fig. 5B), thereby supporting the involvement of a silencing mechanism similar to that described for *GABRQ* and *GABRA3* (Fig. 4D). Interestingly, eight among the overlapped hypermethylated genes (*WT1, PAX6, GNAS, EPB41L1, CSMD1, CPEB1, RERG*, and *SMAD6*) were previously reported to exhibit tumor suppressive functions.

**Figure 5.**
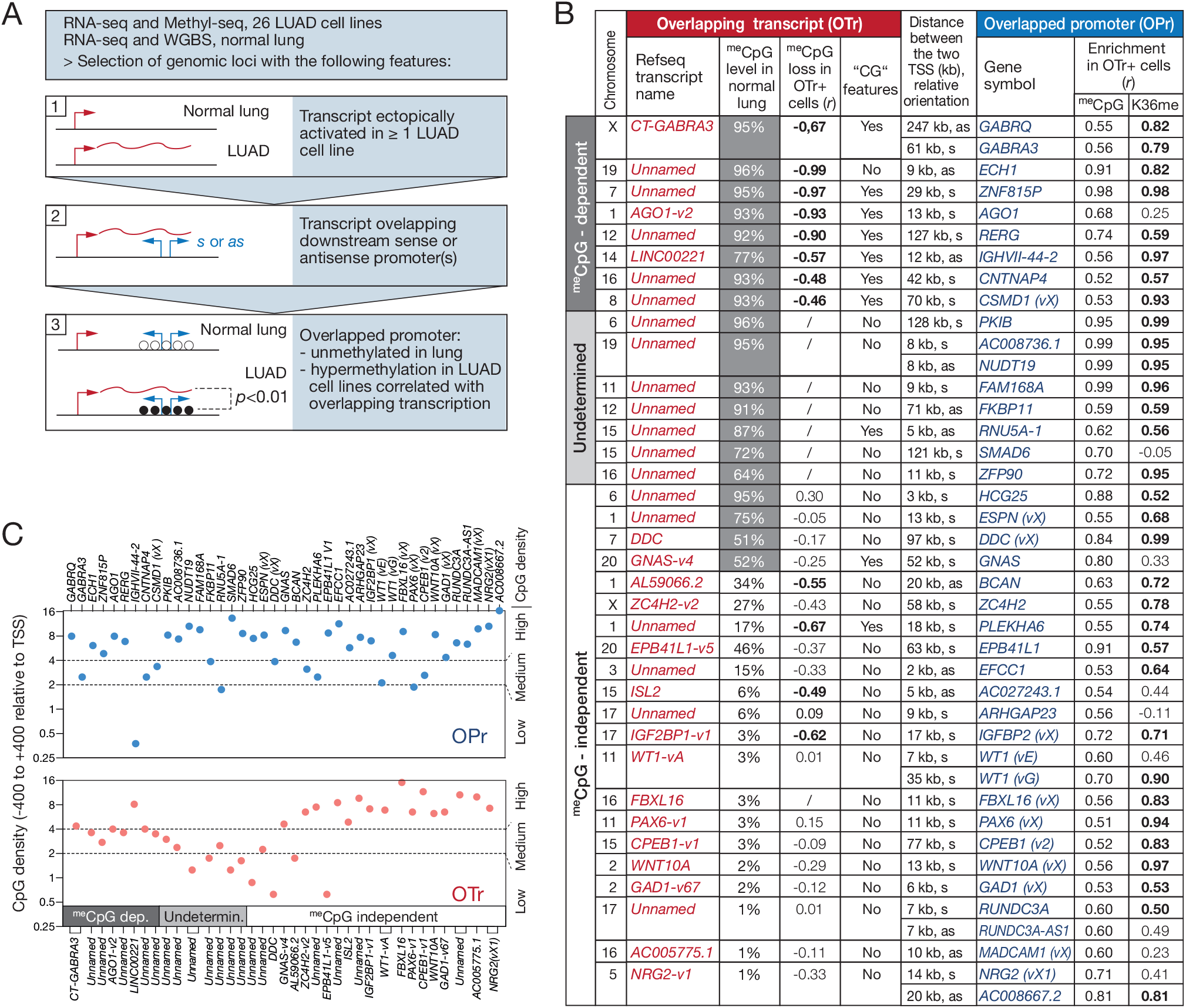
Identification of other genomic loci displaying correlation between DNA hypermethylation of an overlapped promoter (OPr) and activation of an overlapping transcript (OTr) in LUAD cell lines. (*A*) Flowchart of the bioinformatics selection procedure. Broken arrows indicate positions of transcription start sites, wavy lines correspond to overlapping transcription, and empty or filled circles represent unmethylated or methylated CpGs, respectively. (*B*) Features of selected genomic loci. For OPrs, the table provides: distance from TSS of OTr; gene symbol (with transcript variant according to NCBI RefSeq descriptions, or indicated as vX when the variant was undescribed); Pearson coefficient of correlation between OTr activation and promoter CpG methylation; Pearson coefficient of correlation between OTr activation and promoter H3K36me3 (coefficient in bold if adjusted *p-*value < 0.05). For OTrs: Name of transcript or transcript variant, or “unnamed” if not described in NCBI RefSeq; CpG methylation level of 5’-region in normal lung; Pearson coefficient of correlation between transcriptional activation and 5’-region CpG methylation level (“/” when methylation data were unavailable, coefficient in bold if adjusted *p-*value < 0.05); cancer-germline (“CG”) features, i.e. OTr is specifically expressed and its 5’-region demethylated in testis. OTrs were categorized according to their predicted dependency (^me^CpG-dependent) or independency (^me^CpG-independent) on DNA demethylation for transcriptional activation in LUAD cells (see text for criteria); several OTrs were categorized “Undetermined” because methylation data in LUAD cell lines were lacking. (*C*) The number of CpGs were calculated in the genomic segments located -400 to +400 relative to the TSS of OPrs and OTrs. Results are expressed as number of CpGs per 100 bp, and are plotted on a log2 scale. Sequences were categorized according to CpG density: (>4 CpG/100 bp: High, 2 to 4 CpG/100 bp: Medium, <2 CpG/100 bp: Low).

### DNA demethylation accounts for the ectopic activation of several overlapping transcripts

We next searched to determine if DNA hypomethylation accounted for activation of the overlapping transcripts in the genomic loci we selected. To this end, we first exploited bisulfite-seq data from normal human tissues in order to sort out OTrs that have their promoter initially methylated in normal lung (^me^CpG ≥50%, Fig. 5B). In addition, we examined Methyl-seq data from DBTSS, in order to identify OTrs that show significant association between activation and promoter demethylation among the 26 LUAD cell lines (Fig. 5B). Seven OTr genes (besides *CT-GABRA3*) fulfilled the two criteria, and were therefore considered as being DNA methylation dependent. Importantly, 6 out of these 7 genes displayed typical “cancer-germline” features, i.e. preferential expression and promoter demethylation in testis (Fig. 5B, supplementary Fig. S5). Moreover, 6 of these OTr genes contained an intermediate density of CpGs within their 5’ region (Fig. 5C), a recognized characteristic of DNA methylation-regulated gene promoters [30]. For 7 other OTr genes, high promoter methylation was observed in normal lung, but Methyl-seq data in LUAD cell lines were lacking. Dependency on DNA methylation could therefore not be determined for these genes. The remaining 17 OTr genes were considered to be regulated by mechanisms not involving DNA methylation (Fig. 5B). Together, our selection procedure in LUAD cell lines led to the identification of 7 genes besides *GABRQ* and *GABRA3* (*ECH1, ZNF815P, AGO1, RERG, IGHVII-44-2, CNTNAP4, CSMD1*) that become hypermethylated in lung tumor cells most likely through a process of DNA hypomethylation-induced overlapping transcription. Hypermethylation of other genes were also found to be associated with transcriptional overlap, but in these cases activation of the overlapping transcript did not appear to be due to promoter DNA demethylation.

### DNA hypomethylation-induced transcriptional overlap is linked with promoter hypermethylation of *PTPRO* and *RERG* tumor suppressor genes

We chose to further investigate the *RERG* locus on chromosome 12, as it turned out that the OTr in this region overlaps not only one, but two genes with previously reported tumor suppressor functions: *RERG* (RAS Like Estrogen Regulated Growth Inhibitor), a negative regulator of the RAS/MAPK pathway and inhibitor of cell proliferation and tumor formation [31, 32]; and *PTPRO* (Protein Tyrosine Phosphatase Receptor Type O), a phosphatase that counteracts the activity of tyrosine kinases, and modulates cell cycle progression and apoptosis [33, 34].

Examination of RNA-seq data with the Splice Junctions analysis tool of the Integrative Genome Viewer (IGV) confirmed the presence of a transcript overlapping *PTPRO* and *RERG* promoters in testis and several LUAD cell lines (Fig. 6A). The OTr was therefore named *CT-RERG* (Cancer-Testis *RERG*). RT-PCR experiments and RNA-seq data in healthy tissues revealed that *CT-RERG* is expressed not only in testis but also in placenta (Fig. 6B, and supplementary Fig. S5), a feature shared by several CG genes. (Fig. 6B). Despite the presence of the entire *RERG* open reading frame in the *CT-RERG* mRNA, this transcript variant appeared as a poor substrate of RERG protein translation, probably due to the presence of short upstream open reading in the specific 5’ exons (supplementary Fig. S6).

**Figure 6.**
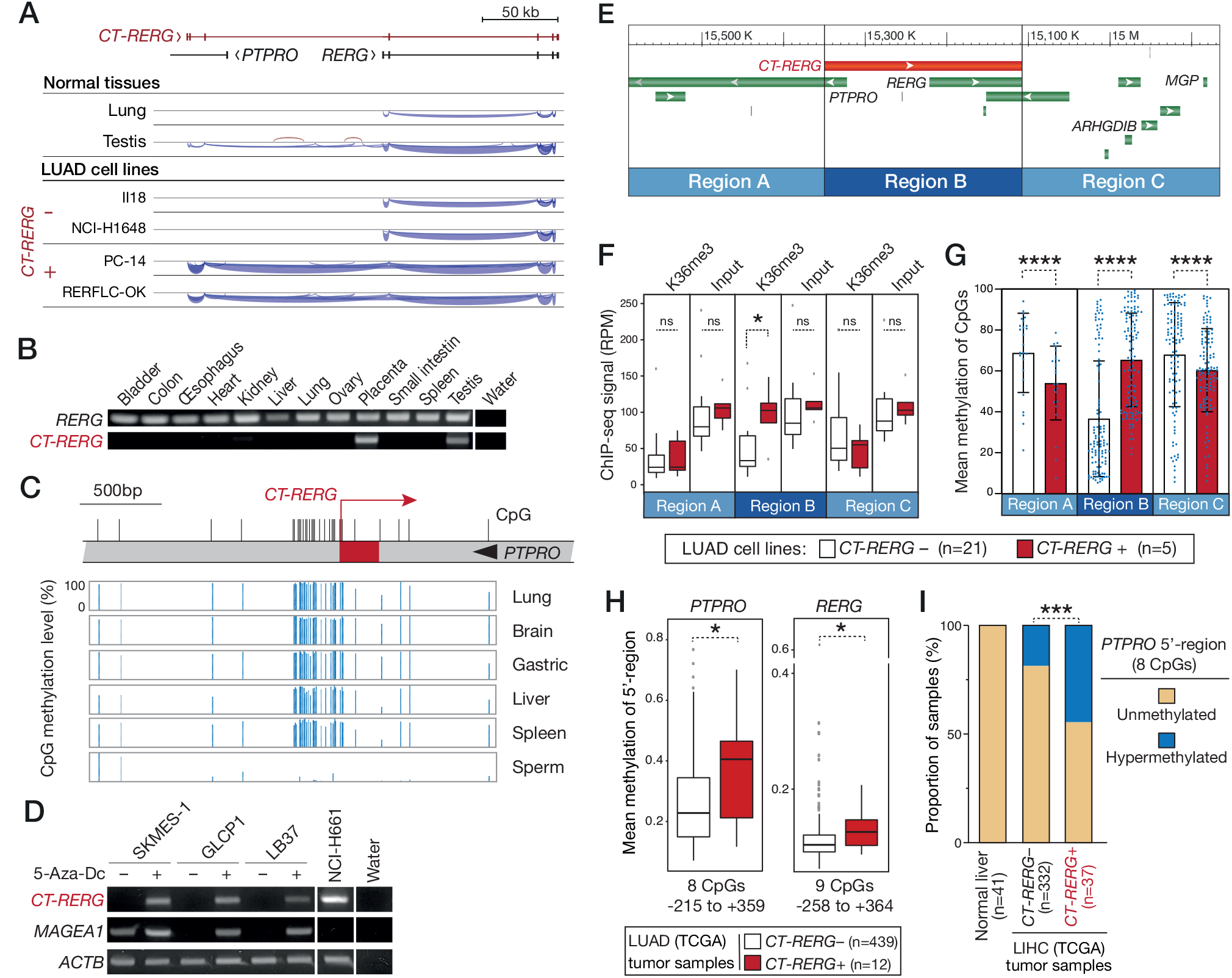
DNA hypomethylation-induced transcriptional overlap is associated with DNA hypermethylation of *PTPRO* and *RERG* tumor suppressor genes. (*A*) RNA-seq data from indicated samples were analyzed with IGV. Exon-intron structures are depicted on the top. (*B*) RT-PCR analyses in various normal tissues were performed with primers specific for either *RERG* of *CT-RERG* transcripts. (*C*) Bisulfite-seq data (NIH Roadmap epigenomics) revealing the methylation level (histograms) of CpG sites (vertical bars) surrounding the *CT-RERG* transcription start (broken arrow) in normal tissues. (*D*) *CT-RERG*-negative lung tumor cell lines were cultured with (−) or without (+) the DNA methylation inhibitor 5-azadC, and RT-PCR experiments were performed to test induction of *CT-RERG* mRNA. NCI-H661 lung tumor cell line was used as a positive control of *CT-RERG* expression. *ACTB* served as an internal control, and *MAGEA1* as a control of 5-azadC induction. (*E*) Three 240 kb genomic regions were defined for subsequent analyses: *CT-RERG* transcription unit (B) and the two neighboring segments (A and C). (*F*) Mean amounts of ChIP-seq reads for H3K36me3 or control input in genomic regions A, B and C were compared in LUAD cell lines that do or do not express *CT-RERG*. * Mann-Whitney test, *p*-value <0.05. (*G*) Mean methylation levels (%, Methyl-seq data) of all CpGs contained in each of the three genomic regions were compared in *CT-RERG*-positive and -negative LUAD cell lines. **** Mann-Whitney test, *p*-value ≤0.0001. (*H*) Mean methylation levels of all CpG sites embedded within the 5’region of either the *PTPRO* (n=8, -215 to +359 relative to TSS) or *RERG* (n=9, -258 to +364) were compared in *CT-RERG*-and *CT-RERG*+ LUAD tumor samples (TCGA). * Mann-Whitney test, *p*-value <0.05. (*I*) The proportion of samples with *PTPRO* hypermethylation was determined in TCGA samples of hepatocellular carcinoma (n=369) grouped according to the *CT-RERG* expression status. Considering that the mean CpG methylation values within the *PTPRO* 5’-region was 0.08 (SD±0.014) in normal liver tissues (n = 41), the region was considered hypermethylated in tumor samples where the mean CpG methylation value was > 0.2. *** Fisher’s exact test, *p*-value <0.001.

Analysis of bisulfite-seq datasets showed that CpG sites located around the TSS (−/+400 bp) of *CT-RERG* are highly methylated in normal somatic tissues and instead almost completely unmethylated in sperm (Fig. 6C). Experimental induction of *CT-RERG* upon treatment with the DNA demethylating agent 5’-aza-deoxycytidine (5-Aza-dC) was observed in all of three tested tumor cell lines (Fig. 6D), thereby confirming the primary role of DNA methylation in its regulation.

As for *CT-GABRA3*, we observed that transcriptional activation of *CT-RERG* in LUAD cell lines was accompanied by increases in H3K36 tri-methylation over the entire 240 kb-long transcription unit, while neighboring regions remained unaffected (Fig. 6E,F). Consistently, whereas the *CT-RERG* transcription unit (region B) showed increased DNA methylation in expressing LUAD cell lines, neighboring regions A and C displayed instead reduced DNA methylation levels in these cell lines (Fig. 6G). Analyses of the *RERG* locus were further extended to *in vivo* tumor samples. Examination of TCGA datasets demonstrated that *CT-RERG* transcription is significantly correlated with *PTPRO* and *RERG* promoter hypermethylation in lung adenocarcinoma tissues (Fig. 6H). Since *PTPRO* has also been reported to exert a tumor suppressor function in hepatocellular carcinoma [33], we analyzed corresponding TCGA datasets to verify association between *CT-RERG* expression and *PTPRO* hypermethylation in this tumor type. The results confirmed increased frequencies of *PTPRO* hypermethylation in the hepatocellular carcinoma samples that express *CT-RERG* (Fig. 6I). Together these observations confirm that DNA hypomethylation is associated with *CT-RERG* expression, and consequently with an increased propensity for *PTPRO* and *RERG* promoters to become hypermethylated. We noticed however, that a few *CT-RERG*-negative tumor samples nevertheless displayed DNA hypermethylation of *PTPRO* and *RERG* (Fig. 6H and I), thereby suggesting that transcriptional overlap may not be the only mechanism directing epigenetic silencing onto these promoters.

## DISCUSSION

It is currently proposed that DNA hypomethylation contributes to tumor progression by inducing genome instability, and by activating genes with oncogenic potential [12, 35]. Our study now raises the interesting, and paradoxical, possibility that it also favors tumor development by contributing indirectly to the repression of tumor suppressor genes. We found indeed that focal DNA hypomethylation in tumor cells can lead to aberrant activation of transcripts that overlap downstream promoters and trigger their hypermethylation. Our work establishes therefore an unrecognized connection between DNA hypomethylation and DNA hypermethylation in tumors. This epigenetic coupling, however, applies to discrete genomic sites, and is therefore compatible with the accepted notion that genome-wide DNA hypomethylation is not associated at the global level with higher frequencies of DNA hypermethylation events [5]. Tumor-type specific patterns of DNA hypomethylation and hence of overlapping transcript activation, may instead be partly responsible for the selectivity of DNA hypermethylation events that is observed among tumors of different origins [36, 37]. For instance, hypermethylation of *PTPRO* and *RERG* promoters was only occasionally detected in renal carcinoma, a tumor type known to display infrequent hypomethylation and activation of CG genes [11], and in which we found seldom activation of *CT-RERG* (supplementary Fig. S7).

The role of overlapping transcription in directing *de novo* methylation of downstream promoters has been previously documented in normal developmental processes, notably during differentiation of embryonic stem cells, and for the establishment of parental imprinting marks in the germline [16, 19, 38]. Although transcription is an efficient way of inducing *de novo* methylation of downstream CpGs, our analyses showed that overlapped promoters sometimes remained unmethylated. Lack of *GABRQ*/*GABRA3* and *PTPRO*/*RERG* promoter hypermethylation was indeed observed in a fraction of tumor samples that nevertheless produced the overlapping transcripts. Hypermethylation of these promoters was also absent in testicular germ cells, where overlapping transcripts are expressed (supplementary Fig S8). It is therefore likely that, under certain conditions, promoters can resist overlapping transcription-induced DNA hypermethylation. Such a mechanism of resistance was previously reported in differentiating embryonic stem cells, and was correlated with elevated transcriptional activity of the overlapped promoter [16]. Moreover, we hypothesize that overlapped promoters are at higher risk of becoming hypermethylated in tumors that exhibit molecular imbalances favoring *de novo* DNA methylation, for instance through exacerbated activities of DNMT methyltranferases or impaired functioning of TET demethylases [39-42].

Transcripts that overlap *GABRA3* and *RERG* promoters are in sense orientation, and share all coding exons with the corresponding overlapped genes. Our analyses revealed, however, that these overlapping transcripts do not produce the corresponding proteins, possibly due to the presence of short upstream ORFs in the specific 5’ exons (supplementary Fig. S6) and [15]. When activated, these non-coding overlapping transcripts do have therefore the potential to cause loss of function of the overlapped gene. A corollary to this observation is that analyses of transcriptomic data in tumors might in some cases suggest that a gene is activated, when in fact activation pertains to a non-coding overlapping transcript that actually leads to loss of function of the gene. This may partly explain previous observations linking DNA hypermethylation with transcriptional activation [43, 44]. Hence, high-resolution analyses of transcriptomic and methylomic data are required in order clearly understand the links between DNA methylation changes and gene expression in tumors [45].

It is hoped that a better understanding of the processes that underlie epigenetic alterations in tumors will lead to the development of novel tools for the diagnosis and therapy of cancer. In this line, establishing the pattern of expression of overlapping transcripts in tumor samples could serve to predict tumor suppressor genes that are at risk to become hypermethylated. Moreover, epigenetic anti-cancer therapies aiming at reactivating silenced tumor suppressor genes might benefit from the knowledge that some of these genes owe their hypermethylated status to a process of transcriptional overlap, and therefore to the specific contribution of druggable chromatin regulators, such as modifiers and readers of H3K36me3 marks.

## Acknowledgements

This work was supported by grants from the D.G. Higher Education and Scientific Research of the French Community of Belgium (Action de Recherches Concertées) and from the Fonds special de recherche (FSR) of the UCLouvain, Belgium. AL was supported by the de Duve Institute, Brussels, Belgium. AVT was a recipient of a Télévie grant from the FRS-FNRS, Belgium [#7.4581.13]. AD received a fellowship from FRS-FNRS-FRIA, Belgium [#1.E008.19]. Funding for open access charge: de Duve Institute, Brussels, Belgium.

## Author’s contributions

J.S.F., A.L. and C.D.S conceived and designed the experiments: J.S.F., A.V.T., A.L. performed the experiments. J.S.F., A.L., A.D. and C.D.S. analysed the data. J.F.S. and C.D.S. wrote the paper.

