## Supplemental methods, figures and tables for "Transcriptional overlap links DNA hypomethylation with DNA hypermethylation at adjacent promoters in cancer"

**This PDF file includes:**

Supplementary methods
Figures S1 to S8
Tables S1 to S3
Supplementary References

### Supplementary Methods

**Processing of public RNA-seq raw data.**
*1) Read alignment*: Fastq files from technical replicates, when available, were summed to increase sequencing depth. FastQC v0.11.8 was used for read quality control (1). Low-quality reads were discarded using Trimmomatics v0.38 (2). Resulting files were mapped to the genome GRCh38 using HISAT2 v2.1.0 with default parameters (3), and were converted to BAM files and indexed using Samtools v1.6 (4). Read alignments corresponding to splice junctions were visualized using the Integrative Genomics Viewer v2.3.68 (IGV) (5). *2) De novo transcriptome assembly and quantification of full-length referenced and unreferenced transcripts:* StringTie v1.3.4 assembling was applied to BAM files of the 26 LUAD cell lines, normal lung, and testis tissues (using -G option for GENCODE v27), and resulting GTF files were merged (--merge option) to produce a unified non-redundant set of transcripts (6). This merged GTF file was used for a re-quantification step in LUAD cell lines, lung, and testis. To this end, the BAM file for each sample was submitted to StringTie assembling (-e -B -G parameters), using the merged GTF as the reference transcriptome. Transcript expression levels are expressed as Transcripts Per Million (TPM).

**Bioinformatics workflow for the identification of overlapped promoter hypermethylation**.
*1) Pooling of transcripts originating from the same promoter in LUAD cell lines:* Transcripts originating from a TSS located less than 50 bp from each other were pooled for quantification, i.e. their expression level corresponded to the sum of individual transcript counts. For such pooled transcripts, the TSS most proximal to the 5' end was used as reference. *2) Selection of activated transcripts and potential overlapped promoters in LUAD cell lines:* Transcripts that are initially not expressed in normal lung (TPM < 0.1 in normal lung) were retained, and among these, those showing expression (> 10 TPM) in at least 1 to maximum 20 of the 26 LUAD cell lines were selected. Finally, transcripts overlapping another TSS, whatever its orientation, were identified. *3) Correlation between overlapping transcript expression and overlapped promoter methylation:* For each of the potential overlapped promoters, mean methylation levels in normal lung and LUAD cell lines (TSS +/- 400bp; promoters with less 3 CpGs within this range were discarded) was computed. Inclusion criteria for potentially hypermethylated overlapped promoters were: mean methylation level in normal lung < 40%; methylation data available in at least 10/26 LUAD cell lines; not all cell lines displaying > 40% methylation. Overlapped promoters that passed this sorting were then examined for potential correlation between their DNA methylation level and expression of the overlapping transcript in LUAD cell lines. Genomic loci displaying a correlation coefficient > 0.3 (*p*-value < 0.01), were selected. For overlapping transcripts corresponding to pooled transcripts (see step 1), the expression level was calculated on the basis of only transcripts variants that actually do overlap the downstream promoter. *4) Manual curation of overlapping transcript/overlapped promoter pairs.* To verify accuracy of TSS positions and transcript structures inferred from our transcriptome *de novo* assembly (see above), BAM files of LUAD cell lines were visualized using IGV (5). This led to a final selection of genomic loci, in which overlapping transcription was confirmed, and where position of the TSS of both the overlapping transcript and overlapped promoter could be validated.

### Supplementary Figures

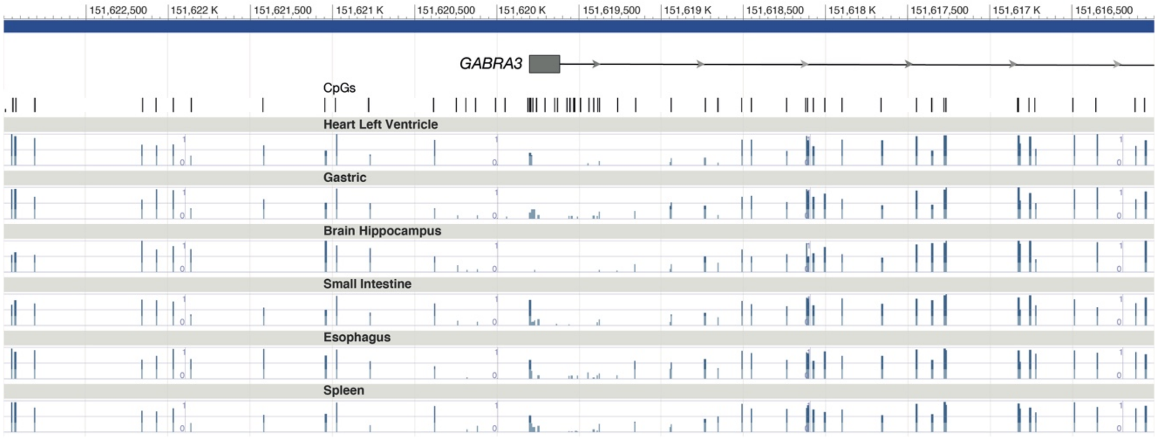


**Fig. S1.** CpGs within the 5'-region of *GABRA3* are mostly unmethylated in normal tissues. Bisulfite-seq data from different normal human tissues (all from males) were visualized with the NIH Roadmap epigenomics track viewer: histograms depict CpG methylation levels (fraction methylated). Position of CpG sites and of exon 1 of *GABRA3* are depicted above.

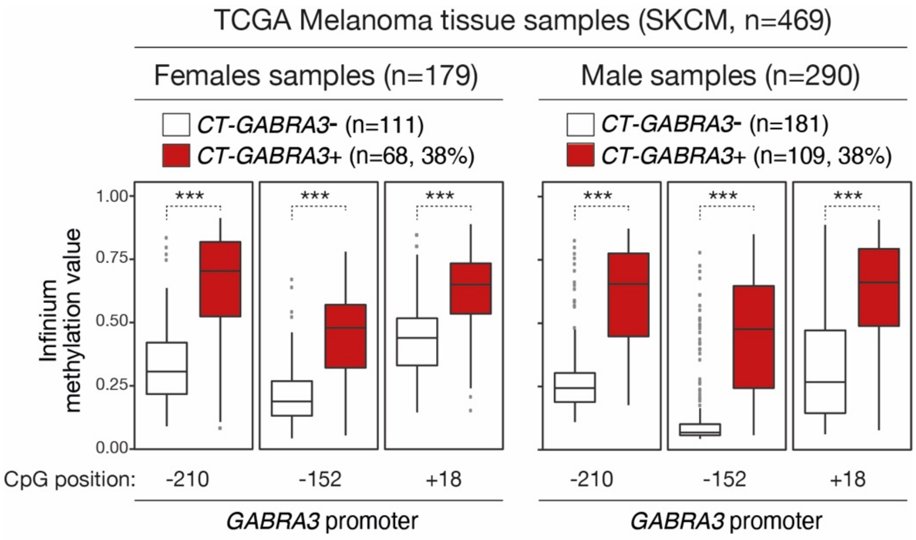


**Fig. S2.** Hypo/hypermethylation in the *GABRA3* locus is not affected by gender. Melanoma tissue samples from the TCGA were grouped according to gender and to the *CT-GABRA3* expression status (inferred from RNA-seq data). The frequency of activation of *CT-GABRA3* was identical in melanoma samples of male and female origin (38%). The methylation level of three CpG sites embedded within the *GABRA3* 5'-region (position relative to TSS) were determined in the different groups of melanoma samples through the analysis of Infinium methylation data (probe intensity ratio). Samples of both male and female showed significant association between *CT-GABRA3* activation and hypermethylation of CpG sites within the *GABRA3* promoter region. *** Welch's t-test, adjusted p-value <0.001.

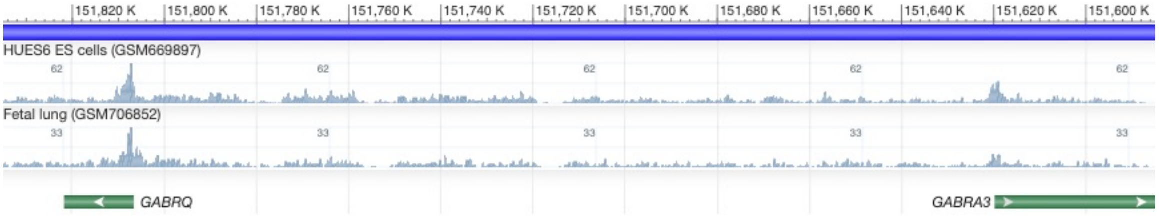


**Fig. S3.** *GABRA3* and *GABRQ* promoters are pre-marked with H3K27me3. H3K27me3 ChIP-seq data for indicated human samples were visualized with the NIH Roadmap epigenomics track viewer: histograms depict H3K27me3 enrichment. Chromosome (X) coordinates refer to the GRCh37.p13 genome assembly release.

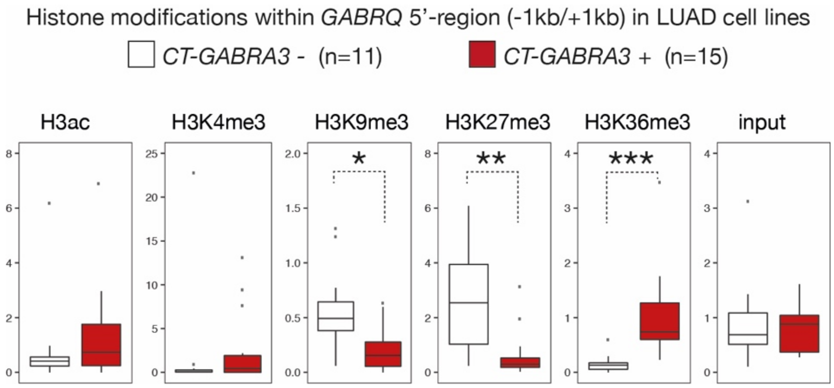


**Fig. S4.** *CT-GABRA3* transcription correlates with decreased H3K9me3/H3K27me3, and increased H3K36me3 within the 5'-region of *GABRQ*. (A) ChIP-seq results for each histone modification were analyzed for the *GABRQ* 5'-region (TSS +/- 1kb) in LUAD cell lines that do or do not express *CT-GABRA3*. Analysis was carried out as described in the material and methods section. *, **, and *** Mann Whitney test, adjusted *p*-value <0.05, <0.01, and <0.001, respectively.

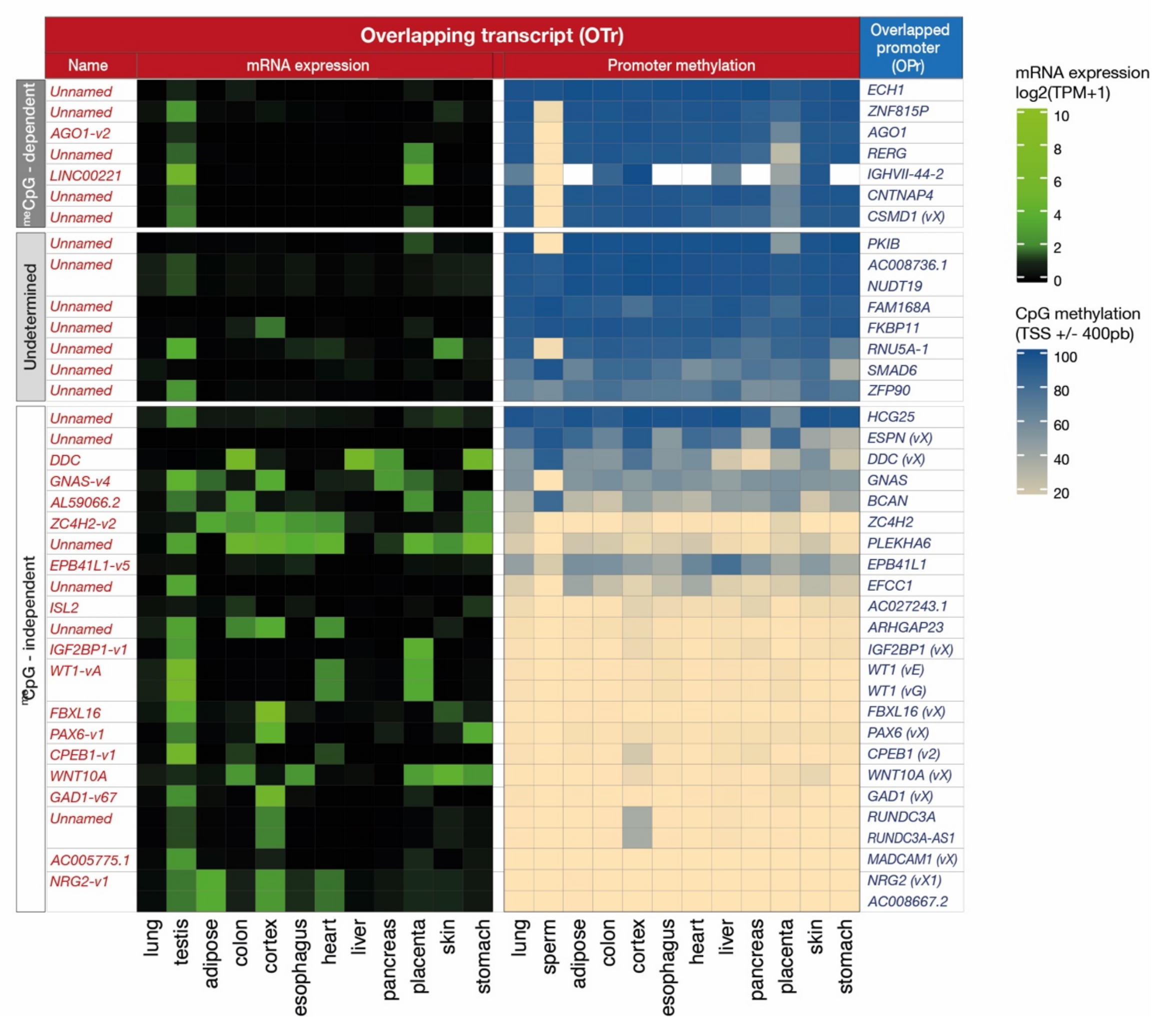


**Fig. S5.** Patterns of mRNA expression and mean promoter DNA methylation (TSS +/- 400bp) of overlapping transcripts (OTrs) in normal human tissues. Public raw RNA-seq data of normal tissues were downloaded from SRA (supplemental table S3) and were processed using the same pipeline as for lung and testis described in the material and methods section. Transcripts originating from the same TSS for a specific locus were summed, as described in the material and methods section. For WGBS, fastq files of normal cortex tissue were downloaded from SRA (supplemental table S3). Read quality control was performed using Trim-Galore v0.5.0 and FastQC v0.11.8. Technical replicates were summed. Read alignment and methylation calling were performed using Bismark v0.20.0. BAM file of liver (supplemental table S3) was downloaded from ENCODE. Methylation calling was performed using Bismark v0.20.0. For all other normal tissues, normalized hg38 data were downloaded from ENCODE (supplemental table S3).

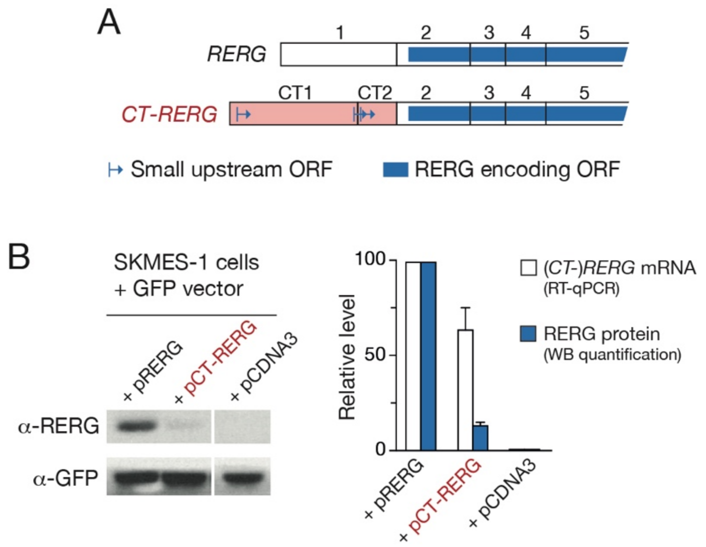


**Fig. S6.** Short upstream open reading frames (uORFs) in *CT-RERG* mRNA inhibit translation of the RERG protein. (*A*) Examination of the first two specific exons of *CT-RERG*, however, revealed the presence of three short upstream open reading frames (uORFs). Several reports have shown that uORFs can be associated with reduced protein expression by inhibiting translation initiation from the main downstream ORF (7,8). (*B*) We therefore compared the ability of *CT-RERG* and *RERG* transcripts to lead to production of the RERG protein. To this end, expression vectors carrying the corresponding cDNAs (pRERG and pCT-RERG), as well as the empty vector (pCDNA3), were transfected together with a GFP-expressing vector into the SKMES-1 lung tumor cell line. Western blotting (WB), performed at day 3 after transfection, examined the presence of the RERG protein and GFP as an internal control. Three experimental replicates were performed (one illustrative WB is shown). Band intensities in WBs were quantified with ImageJ. From the same transfected cell groups, we purified DNAse-treated RNA to check (*CT-*)*RERG* relative expression levels by RT-qPCR analysis (RT-minus subtraction and *ACTB* normalization were applied). The results revealed a marked decrease of RERG protein levels in p*CT-RERG* transfectants (14%), when compared with pRERG transfectants (taken as 100% reference). *CT-RERG* mRNA levels in p*CT-RERG* transfectants were only moderately lower (63%) than *RERG* mRNA level in pRERG transfectants. These data reveal therefore marked inhibition of RERG protein translation in *CT-RERG* transcripts. *Detailed protocol: RERG* and *CT-RERG* 5'-UTRs were amplified by PCR using the high fidelity PrimeStar HS DNA polymerase (Takara) with primers carrying a 5' overhang with a restriction site for *Hind*III (forward primer) and *Xho*I (reverse primer) (Table S2). PCR fragments were cloned into pcDNA3.1 expression vector (Invitrogen), and verified by sequencing. pTM624 GFP expressing vector was kindly provided by Thomas Michiels (de Duve Institute). SKMES-1 cells were plated at 150.000 cells/well of a 6w plate the day before transfection. Cells were transfected with 1.5 µg of plasmid pRERG, pCT-RERG or empty pcDNA3.1 and 0.5 µg of pTM624 using 6 µl Viafect transfection reagent (Promega) according to manufacturer's instructions. RNA (treated with Turbo DNAse, Invitrogen) and proteins were harvested 3 days post-transfection. For WB analysis, both membranes for RERG and GFP detection were blocked in 5% milk in TBS-Tween 0.1%. For detection of RERG protein, rabbit polyclonal anti-RERG antibody (10687-1-AP, Sanbio) was used at a final dilution of 1/5000 in 5% milk in TBS-Tween 0.1%. HRP-conjugated goat anti-rabbit secondary antibody (ADI-SAB-300-J, Enzo Life Sciences) antibody was used at 1/10000 in TBS-Tween 0.1%. For GFP protein detection, we used goat polyclonal anti-GFP antibody (ab6673, Abcam) at a final dilution of 1/5000 in 5% milk in TBS-Tween 0.1%. HRP-conjugated donkey anti-goat (PA1-28664, Life Technologies) was used at a final dilution 1/5000 in TBS-Tween 0.1%.

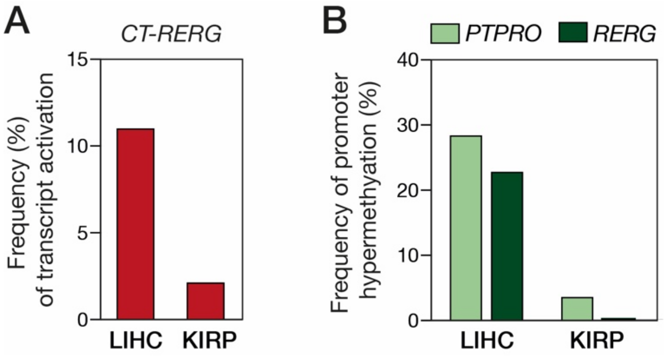


**Fig. S7.** Renal carcinoma shows lower frequency of both transcriptional activation of *CT-RERG* and hypermethylation of *PTPRO* and *RERG* downstream promoters. (*A*) The frequency of activation of *CT-RERG* transcription (TPM>1) was evaluated in liver hepatocarcinoma (LIHC, n=369) and renal papillary cell carcinoma (KIRP, n=273) through the analysis of RNA-seq data from the TCGA. (*B*) The frequency of hypermethylation of *PTPRO* and *RERG* promoters in these same tumor samples were evaluated through the analysis of Infinium methylation data from the TCGA (probe intensity ratio > 0.2, as explained in Fig.6H-I).

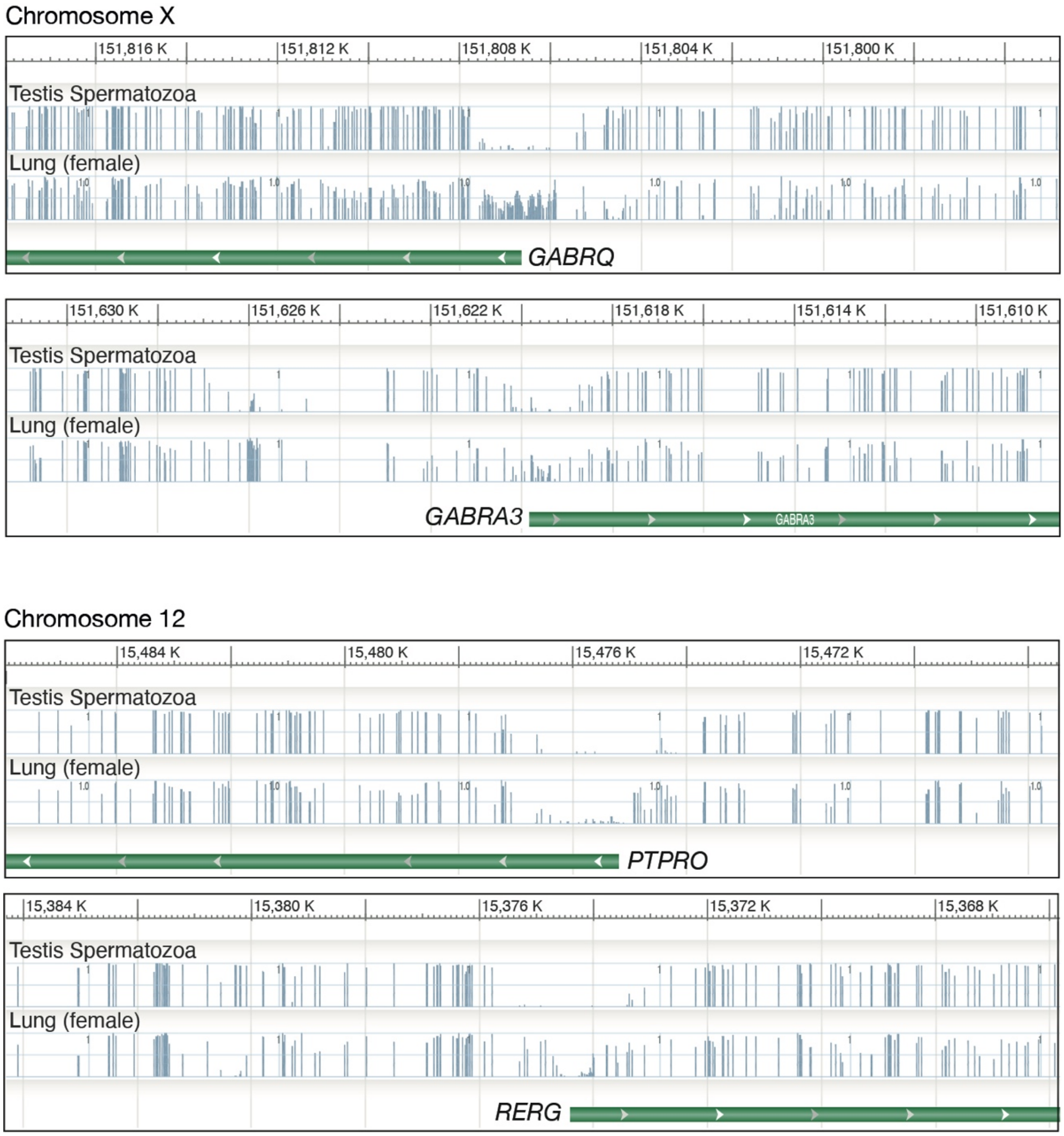


**Fig. S8.** Lack of hypermethylation of *GABRQ*/*GABRA3* and *PTPRO*/*RERG* promoters in testicular germ cells. Bisulfite-seq data from human testicular spermatozoa and normal lung were visualized with the NIH Roadmap epigenomics track viewer: histograms depict CpG methylation levels (fraction methylated). TSS position and orientation (green bar and arrowhead) of the four genes analyzed are indicated. Partial methylation of *GABRQ* and *GABRA3* gene promoters in the lung is attributable to the presence of an inactive X chromosome, as the tissue was obtained from a female individual.

### Supplementary Tables

**Table S1.** Transcript references and TSS positions of overlapping transcripts (OTr) and overlapped promoters (OPr) identified in this study.

| | Overlapping transcripts (OTr) | | | | | Overlapped promoter (Opr) | | | |
|---|---|---|---|---|---|---|---|---|---|
| Chromosome | NCBI Refseq description | Corresponding reference(s) in Ensembl (ENST) or in-house generated GTF file (MSTRG) | Position of TSS | Strand | | Gene symbol (variant) | Corresponding ENST or MSTRG | Position of TSS | Strand |
| chrX | CT-GABRA3 | MSTRG.42953.5, MSTRG.42953.6, MSTRG.42953.8, MSTRG.42953.4, MSTRG.42953.3, MSTRG.42953.15, MSTRG.42953.7, MSTRG.42953.13, MSTRG.42953.14, MSTRG.42953.12 | 152698719 | - | -> | GABRQ | ENST00000598523.2 | 152638185 | + |
| | | | | | | GABRA3 | ENST00000370314.8 | 152451315 | - |
| chr19 | Unnamed | MSTRG.20900.2 | 38823300 | + | -> | ECH1 | ENST00000221418.8 | 38832005 | - |
| chr7 | Unnamed | MSTRG.35963.2 | 5794482 | + | -> | ZNF815P | ENST00000421890.5 | 5823160 | + |
| chr1 | AGO1-v2 | ENST00000373206.5 | 35869808 | + | -> | AGO1 | ENST00000373204.5 | 35883209 | + |
| chr12 | Unnamed (CT-RERG) | MSTRG.8305.2 | 15348675 | - | -> | RERG | ENST00000256953.6 | 15221417 | - |
| chr14 | LINC00221 | ENST00000603633.2 | 106482439 | + | -> | IGHVII-44-2 | ENST00000517728.1 | 106494383 | - |
| chr16 | Unnamed | MSTRG.16001.1 | 76235202 | + | -> | CNTNAP4 | ENST00000611870.4 | 76277278 | + |
| chr8 | Unnamed | MSTRG.37947.3 | 3183689 | - | -> | CSMD1 (vX) | ENST00000520561.1 | 3113363 | - |
| chr6 | Unnamed | MSTRG.35314.8 | 122482079 | + | -> | PKIB | ENST00000368452.6 | 122610232 | + |
| chr19 | Unnamed | MSTRG.20651.2 | 32700158 | - | -> | AC008736.1 | ENST00000592431.1 | 32691750 | - |
| chr19 | Unnamed | MSTRG.20651.2 | 32700158 | - | -> | NUDT19 | ENST00000397061.3 | 32691961 | + |
| chr11 | Unnamed | MSTRG.7040.15 | 73607387 | - | -> | FAM168A | ENST00000064778.8 | 73598189 | - |
| chr12 | Unnamed | MSTRG.8692.1 | 48854530 | + | -> | FKBP11 | ENST00000550765.5 | 48925827 | - |
| chr15 | Unnamed | ENST00000558873.1 | 65300618 | - | -> | RNU5A-1 | ENST00000362698.1 | 65296051 | + |
| chr15 | Unnamed | MSTRG.13615.1 | 66582190 | + | -> | SMAD6 | ENST00000557916.5 | 66703259 | + |
| chr16 | Unnamed | MSTRG.15775.2 | 68527848 | + | -> | ZFP90 V1 | ENST00000563169.6 | 68539284 | + |
| chr6 | Unnamed | MSTRG.34416.1 | 33246857 | + | -> | HCG25 | ENST00000450514.5 | 33249534 | + |
| chr1 | Unnamed | MSTRG.191.7 | 6427871 | + | -> | ESPN (vX) | ENST00000418286.1 | 6440378 | + |
| chr7 | DDC | ENST00000444124.6, MSTRG.36458.11 | 50565457 | - | -> | DDC (vX) | MSTRG.36458.12 | 50468168 | - |
| chr20 | GNAS-v4 | ENST00000371075.7 | 58839718 | + | -> | GNAS | ENST00000603546.1 | 58892333 | + |
| chr1 | AL590666.3 | ENST00000448869.1 | 156661424 | - | -> | BCAN | ENST00000329117.9 | 156641948 | + |
| chrX | ZC4H2-v2 | ENST00000337990.2 | 65034713 | - | -> | ZC4H2 | ENST00000374839.7 | 64976484 | - |
| chr1 | Unnamed | MSTRG.3292.8 | 204377853 | - | -> | PLEKHA6 | ENST00000272203.7 | 204359916 | - |
| chr20 | EPB41L1-v5 | ENST00000202028.9 | 36091504 | + | -> | EPB41L1 | ENST00000338074.7 | 36154740 | + |
| chr3 | Unnamed | MSTRG.28969.12 | 129003864 | - | -> | EFCC1 | ENST00000436022.2 | 129001629 | + |
| chr15 | ISL2 | ENST00000290759.8 | 76336724 | + | -> | AC027243.1 | ENST00000559539.1 | 76342063 | - |
| chr17 | Unnamed | MSTRG.17424.1 | 38419280 | + | -> | ARHGAP23 | ENST00000622683.4 | 38428418 | + |
| chr17 | IGF2BP1-v1 | ENST00000290341.7 | 48997171 | + | -> | IGF2BP1 (vX) | ENST00000499130.6 | 49013917 | + |
| chr11 | WT1-vA | ENST00000332351.7 | 32435630 | - | -> | WT1 (vE) | ENST00000527775.1 | 32429029 | - |
| | | | | | | WT1 (vG) | ENST00000526685.1 | 32400615 | - |
| chr16 | FBXL16 | ENST00000397621.5 | 705829 | - | -> | FBXL16 (vX) | ENST00000562585.1 | 695263 | - |
| chr11 | PAX6-v1 | ENST00000640335.1, ENST00000638278.1, ENST00000379107.7, ENST00000379109.7, ENST00000640617.1, ENST00000640684.1, ENST00000638250.1, ENST00000639394.1, ENST00000640242.1, ENST00000639054.1, ENST00000527769.5, ENST00000640613.1, ENST00000471303.6, ENST00000606377.6 | 31813131 | - | -> | PAX6 (vX) | ENST00000640735.1 | 31801992 | - |
| chr15 | CPEB1-v1 | ENST00000615198.4 | 82647977 | - | -> | CPEB1 (v2) | ENST00000562833.2 | 82571447 | - |
| chr2 | WNT10A | ENST00000258411.7 | 218880363 | + | -> | WNT10A (vX) | ENST00000489887.1 | 218893221 | + |
| chr2 | GAD1-v67 | ENST00000358196.8 | 170816114 | + | -> | GAD1 (vX) | ENST00000486850.1 | 170822087 | + |
| chr17 | Unnamed | MSTRG.17685.2, ENST00000588097.5, MSTRG.17685.5 | 44315335 | - | -> | RUNDC3A | ENST00000590834.5 | 44308413 | + |
| | | | | | | RUNDC3A-AS1 | ENST00000586388.1 | 44308393 | - |
| chr19 | AC005775.1 | ENST00000592413.2 | 507833 | - | -> | MADCAM1 (vX) | ENST00000622449.2 | 497817 | + |
| chr5 | NRG2-v1 | ENST00000361474.5 | 140043299 | - | -> | NRG2 (vX1) | ENST00000340391.7 | 139904397 | - |
| | | | | | | AC008667.2 | ENST00000504413.1 | 139848290 | + |

A merged GTF file was generated following *de novo* assembling with StringTie of RNA-seq alignments derived from LUAD cell lines and normal human tissues. Besides transcripts that had been previously described in Ensembl (ENST), novel undescribed transcripts (MSTRG) were identified. For some OTrs, multiple splice variants, all starting from the same promoter, were detected.

**Table S2.** List of primers and PCR conditions.

| | Forward primer (5'-3') | Reverse primer (5'-3') | Additional information |
|---|---|---|---|
| **PCR** | | | |
| *ACTINE-β (ACTB)* | CCCTGGACTTCGAGCAAGAGAT | AAGGTAGTTTCGTGGATGCCACA | DreamTaq (20 cycles / Annealing : 60°C and 30 sec / Elongation : 72°C and 30 sec) |
| *MAGEA1* | GCCGAAGGAACCTGACC | ACTGGGTTGCCTCTGTCG | DreamTaq (20 cycles / Annealing : 60°C and 30 sec / Elongation : 72°C and 30 sec) |
| *RERG* | CTCGCAAACGCAACCTGAA | GAAATCTCACTACAAGAGCTGACTT | DreamTaq (35 cycles / Annealing : 58°C and 30 sec / Elongation : 72°C and 30 sec) |
| *CT-RERG* | TCCTTCACTGTTGTTGGCAGAA | GAAATCTCACTACAAGAGCTGACTT | DreamTaq ( 37 cycles / Annealing : 58°C and 30 sec / Elongation : 72°C and 30 sec) |
| **qPCR** | | | |
| *(CT-)RERG* | GAAGTGCTGCCACTTAAGAACA | GACGCACCTCTCGACACAAT | KAPA SYBR FAST ( Annealing /Elongation : 60°C and 30 sec) |
| **Bisulfite-converted DNA** | | | |
| *CT-GABRA3* first PCR | TYGATTTTTATTTAGGTAGAATTT | TAAAATAATAACRACCCAACCTAA | DreamTaq (35 cycles / Annealing : 55°C and 30 sec / Elongation : 72°C and 30 sec) |
| *CT-GABRA3* second PCR | ATTTAGGTAGAATTTAGTTTTAT | CCCTACRAAATAACCCAAA | DreamTaq (35 cycles / Annealing : 55°C and 30 sec / Elongation : 72°C and 30 sec) |
| *GABRA3* first PCR | AGTGGTGAATTTAAAGTTAGTAAAGG | CCCCAATATCTCCCTACTCAAAT | DreamTaq (35 cycles / Annealing : 55°C and 30 sec / Elongation : 72°C and 30 sec) |
| *GABRA3* second PCR | GATAGAGAGGGAGGGAGGTAGA | CTAAACTTCCACCAACCCCACT | DreamTaq (35 cycles / Annealing : 55°C and 30 sec / Elongation : 72°C and 30 sec) |
| **pcDNA3 Cloning** | | | |
| *RERG* (HindIII & Xho1) | ATGAAAGCTTGCAACTCGCAAACGCAAC | ATGACTCGAGCTAACTACTGATTTTGGTGAGC | PrimeSTAR (37 cycles / Annealing : 62°C and 10 sec / Elongation : 72°C and 1 min) |
| *CT-RERG* (HindIII & Xho1) | ATGAAAGCTTGTTTCACTGTGTTAGCCAG | ATGACTCGAGCTAACTACTGATTTTGGTGAGC | PrimeSTAR (37 cycles / Annealing : 62°C and 10 sec / Elongation : 72°C and 1 min) |

**Table S3.** Transcriptomic and Methylomic datasets used in this study.

| BioSample | Name | Accession number | BioProject | Layout | Instrument | File | Citation |
|---|---|---|---|---|---|---|---|
| RNA-seq datasets | | | | | | | |
| LUAD cell line | A427 | DRR016694 | PRJDB2256 | PAIRED | Illumina HiSeq 2500 | Fastq | DBTSS |
| LUAD cell line | A549 | DRR016695 | PRJDB2256 | PAIRED | Illumina HiSeq 2500 | Fastq | DBTSS |
| LUAD cell line | ABC1 | DRR016696 | PRJDB2256 | PAIRED | Illumina HiSeq 2500 | Fastq | DBTSS |
| LUAD cell line | H1299 | DRR016697 | PRJDB2256 | PAIRED | Illumina HiSeq 2500 | Fastq | DBTSS |
| LUAD cell line | H1437 | DRR016698 | PRJDB2256 | PAIRED | Illumina HiSeq 2500 | Fastq | DBTSS |
| LUAD cell line | H1648 | DRR016699 | PRJDB2256 | PAIRED | Illumina HiSeq 2500 | Fastq | DBTSS |
| LUAD cell line | H1650 | DRR016700 | PRJDB2256 | PAIRED | Illumina HiSeq 2500 | Fastq | DBTSS |
| LUAD cell line | H1703 | DRR016701 | PRJDB2256 | PAIRED | Illumina HiSeq 2500 | Fastq | DBTSS |
| LUAD cell line | H1819 | DRR016702 | PRJDB2256 | PAIRED | Illumina HiSeq 2500 | Fastq | DBTSS |
| LUAD cell line | H1975 | DRR016703 | PRJDB2256 | PAIRED | Illumina HiSeq 2500 | Fastq | DBTSS |
| LUAD cell line | H2126 | DRR016704 | PRJDB2256 | PAIRED | Illumina HiSeq 2500 | Fastq | DBTSS |
| LUAD cell line | H2228 | DRR016705 | PRJDB2256 | PAIRED | Illumina HiSeq 2500 | Fastq | DBTSS |
| LUAD cell line | H2347 | DRR016706 | PRJDB2256 | PAIRED | Illumina HiSeq 2500 | Fastq | DBTSS |
| LUAD cell line | H322 | DRR016707 | PRJDB2256 | PAIRED | Illumina HiSeq 2500 | Fastq | DBTSS |
| LUAD cell line | II18 | DRR016708 | PRJDB2256 | PAIRED | Illumina HiSeq 2500 | Fastq | DBTSS |
| LUAD cell line | LC2ad | DRR016709 | PRJDB2256 | PAIRED | Illumina HiSeq 2500 | Fastq | DBTSS |
| LUAD cell line | PC14 | DRR016710 | PRJDB2256 | PAIRED | Illumina HiSeq 2500 | Fastq | DBTSS |
| LUAD cell line | PC3 | DRR016711 | PRJDB2256 | PAIRED | Illumina HiSeq 2500 | Fastq | DBTSS |
| LUAD cell line | PC7 | DRR016712 | PRJDB2256 | PAIRED | Illumina HiSeq 2500 | Fastq | DBTSS |
| LUAD cell line | PC9 | DRR016713 | PRJDB2256 | PAIRED | Illumina HiSeq 2500 | Fastq | DBTSS |
| LUAD cell line | RERFLC-Ad1 | DRR016714 | PRJDB2256 | PAIRED | Illumina HiSeq 2500 | Fastq | DBTSS |
| LUAD cell line | RERFLC-Ad2 | DRR016715 | PRJDB2256 | PAIRED | Illumina HiSeq 2500 | Fastq | DBTSS |
| LUAD cell line | RERFLC-KJ | DRR016716 | PRJDB2256 | PAIRED | Illumina HiSeq 2500 | Fastq | DBTSS |
| LUAD cell line | RERFLC-MS | DRR016717 | PRJDB2256 | PAIRED | Illumina HiSeq 2500 | Fastq | DBTSS |
| LUAD cell line | RERFLC-OK | DRR016718 | PRJDB2256 | PAIRED | Illumina HiSeq 2500 | Fastq | DBTSS |
| LUAD cell line | VMRC-LCD | DRR016719 | PRJDB2256 | PAIRED | Illumina HiSeq 2500 | Fastq | DBTSS |
| Normal tissue | Lung | SRR577579 SRR577582 | PRJNA34535 | PAIRED | Illumina HiSeq 2000 | Fastq | NIH Roadmap epigenomics |
| Normal tissue | Testis | ERR315352 ERR315415 ERR315492 | PRJEB4337 | PAIRED | Illumina HiSeq 2000 | Fastq | Science for Life Laboratory, Stockholm, Sweden |
| Normal tissue | Esophagus | ERR315411 ERR315398 ERR315489 ERR315434 ERR315472 ERR315362 | PRJEB4337 | PAIRED | Illumina HiSeq 2000 | Fastq | Science for Life Laboratory, Stockholm, Sweden |
| Normal tissue | Colon | ERR315357 ERR315484 ERR315462 ERR315400 ERR315348 ERR315403 | PRJEB4337 | PAIRED | Illumina HiSeq 2000 | Fastq | Science for Life Laboratory, Stockholm, Sweden |
| Normal tissue | Adipose_tissue | ERR315332 ERR315431 ERR315343 ERR315342 ERR315378 | PRJEB4337 | PAIRED | Illumina HiSeq 2000 | Fastq | Science for Life Laboratory, Stockholm, Sweden |
| Normal tissue | Cerebral_cortex | ERR315455 ERR315432 ERR315477 | PRJEB4337 | PAIRED | Illumina HiSeq 2000 | Fastq | Science for Life Laboratory, Stockholm, Sweden |
| Normal tissue | Heart | ERR315384 ERR315328 ERR315356 ERR315367 ERR315413 ERR315435 ERR315389 ERR315331 ERR315430 | PRJEB4337 | PAIRED | Illumina HiSeq 2000 | Fastq | Science for Life Laboratory, Stockholm, Sweden |
| Normal tissue | Liver | ERR315327 ERR315414 ERR315463 ERR315394 ERR315451 | PRJEB4337 | PAIRED | Illumina HiSeq 2000 | Fastq | Science for Life Laboratory, Stockholm, Sweden |
| Normal tissue | Pancreas | ERR315466 ERR315436 ERR315479 ERR315429 | PRJEB4337 | PAIRED | Illumina HiSeq 2000 | Fastq | Science for Life Laboratory, Stockholm, Sweden |
| Normal tissue | Placenta | ERR315374 ERR315478 ERR315336 ERR315375 ERR315476 ERR315377 ERR315399 | PRJEB4337 | PAIRED | Illumina HiSeq 2000 | Fastq | Science for Life Laboratory, Stockholm, Sweden |
| Normal tissue | Skin | ERR315339 ERR315401 ERR315460 ERR315376 ERR315372 ERR315464 | PRJEB4337 | PAIRED | Illumina HiSeq 2000 | Fastq | Science for Life Laboratory, Stockholm, Sweden |
| Normal tissue | Stomach | ERR315379 ERR315369 ERR315467 ERR315485 | PRJEB4337 | PAIRED | Illumina HiSeq 2000 | Fastq | Science for Life Laboratory, Stockholm, Sweden |
| DNA Methylation datasets | | | | | | | |
| LUAD cell line | A427 | bs_data_9606_A427 | DBTSS | PAIRED | Illumina HiSeq 2500 | BED | DBTSS |
| LUAD cell line | A549 | bs_data_9606_A549 | DBTSS | PAIRED | Illumina HiSeq 2500 | BED | DBTSS |
| LUAD cell line | ABC1 | bs_data_9606_ABC1 | DBTSS | PAIRED | Illumina HiSeq 2500 | BED | DBTSS |
| LUAD cell line | H1299 | bs_data_9606_H1299 | DBTSS | PAIRED | Illumina HiSeq 2500 | BED | DBTSS |
| LUAD cell line | H1437 | bs_data_9606_H1437 | DBTSS | PAIRED | Illumina HiSeq 2500 | BED | DBTSS |
| LUAD cell line | H1648 | bs_data_9606_H1648 | DBTSS | PAIRED | Illumina HiSeq 2500 | BED | DBTSS |
| LUAD cell line | H1650 | bs_data_9606_H1650 | DBTSS | PAIRED | Illumina HiSeq 2500 | BED | DBTSS |
| LUAD cell line | H1703 | bs_data_9606_H1703 | DBTSS | PAIRED | Illumina HiSeq 2500 | BED | DBTSS |
| LUAD cell line | H1819 | bs_data_9606_H1819 | DBTSS | PAIRED | Illumina HiSeq 2500 | BED | DBTSS |
| LUAD cell line | H1975 | bs_data_9606_H1975 | DBTSS | PAIRED | Illumina HiSeq 2500 | BED | DBTSS |
| LUAD cell line | H2126 | bs_data_9606_H2126 | DBTSS | PAIRED | Illumina HiSeq 2500 | BED | DBTSS |
| LUAD cell line | H2228 | bs_data_9606_H2228 | DBTSS | PAIRED | Illumina HiSeq 2500 | BED | DBTSS |
| LUAD cell line | H2347 | bs_data_9606_H2347 | DBTSS | PAIRED | Illumina HiSeq 2500 | BED | DBTSS |
| LUAD cell line | H322 | bs_data_9606_H322 | DBTSS | PAIRED | Illumina HiSeq 2500 | BED | DBTSS |
| LUAD cell line | II18 | bs_data_9606_II18 | DBTSS | PAIRED | Illumina HiSeq 2500 | BED | DBTSS |
| LUAD cell line | LC2ad | bs_data_9606_LC2ad | DBTSS | PAIRED | Illumina HiSeq 2500 | BED | DBTSS |
| LUAD cell line | PC14 | bs_data_9606_PC14 | DBTSS | PAIRED | Illumina HiSeq 2500 | BED | DBTSS |
| LUAD cell line | PC3 | bs_data_9606_PC3 | DBTSS | PAIRED | Illumina HiSeq 2500 | BED | DBTSS |
| LUAD cell line | PC7 | bs_data_9606_PC7 | DBTSS | PAIRED | Illumina HiSeq 2500 | BED | DBTSS |
| LUAD cell line | PC9 | bs_data_9606_PC9 | DBTSS | PAIRED | Illumina HiSeq 2500 | BED | DBTSS |
| LUAD cell line | RERFLC-Ad1 | bs_data_9606_RERFLC-Ad1 | DBTSS | PAIRED | Illumina HiSeq 2500 | BED | DBTSS |
| LUAD cell line | RERFLC-Ad2 | bs_data_9606_RERFLC-Ad2 | DBTSS | PAIRED | Illumina HiSeq 2500 | BED | DBTSS |
| LUAD cell line | RERFLC-KJ | bs_data_9606_RERFLC-KJ | DBTSS | PAIRED | Illumina HiSeq 2500 | BED | DBTSS |
| LUAD cell line | RERFLC-MS | bs_data_9606_RERFLC-KJ | DBTSS | PAIRED | Illumina HiSeq 2500 | BED | DBTSS |
| LUAD cell line | RERFLC-OK | bs_data_9606_RERFLC-OK | DBTSS | PAIRED | Illumina HiSeq 2500 | BED | DBTSS |
| LUAD cell line | VMRC-LCD | bs_data_9606_VMRC-LCD | DBTSS | PAIRED | Illumina HiSeq 2500 | BED | DBTSS |
| Normal tissue | Lung | GSM983647 | NIH Roadmap epigenomics | PAIRED | Illumina HiSeq 2000 | BED | NIH Roadmap epigenomics |
| Normal tissue | Sperm | GSM1127119 | NIH Roadmap epigenomics | PAIRED | Illumina HiSeq 2000 | BED | NIH Roadmap epigenomics |
| Normal tissue | Esophagus | ENCFF625GVK | NIH Roadmap epigenomics | PAIRED | Illumina HiSeq 2000 | BED | NIH Roadmap epigenomics |
| Normal tissue | Colon | ENCFF157POM | NIH Roadmap epigenomics | PAIRED | Illumina HiSeq 2000 | BED | NIH Roadmap epigenomics |
| Normal tissue | Adipose_tissue | ENCFF318AMC | NIH Roadmap epigenomics | PAIRED | Illumina HiSeq 2000 | BED | NIH Roadmap epigenomics |
| Normal tissue | Heart | ENCFF536RSX | NIH Roadmap epigenomics | PAIRED | Illumina HiSeq 2000 | BED | NIH Roadmap epigenomics |
| Normal tissue | Pancreas | ENCFF763RUE | NIH Roadmap epigenomics | PAIRED | Illumina HiSeq 2000 | BED | NIH Roadmap epigenomics |
| Normal tissue | Placenta | ENCFF4370KM | NIH Roadmap epigenomics | PAIRED | Illumina HiSeq 2000 | BED | NIH Roadmap epigenomics |
| Normal tissue | Skin | ENCFF219GCQ | NIH Roadmap epigenomics | PAIRED | Illumina HiSeq 2000 | BED | NIH Roadmap epigenomics |
| Normal tissue | Stomach | ENCFF49700 | NIH Roadmap epigenomics | PAIRED | Illumina HiSeq 2000 | BED | NIH Roadmap epigenomics |
| Normal tissue | Liver | ENCFF356KGQ | NIH Roadmap epigenomics | SINGLE | Illumina HiSeq 2500 | BAM | NIH Roadmap epigenomics |
| Normal tissue | Cerebral_cortex | SRR3278486 , SRR3278482 | Johns Hopkins Center for Epigenetics | PAIRED | Illumina HiSeq 2000 | Fastq | Johns Hopkins Center for Epigenetics |
